# Moderate prenatal alcohol exposure modifies sex-specific CRFR1 activity in the central amygdala and anxiety-like behavior in adolescent offspring

**DOI:** 10.1101/2021.09.29.462410

**Authors:** Siara Kate Rouzer, Marvin R. Diaz

## Abstract

Anxiety disorders are highly prevalent among individuals with a history of prenatal alcohol exposure (PAE), and adolescent rodents demonstrate anxiety-like behavior following moderate PAE on Gestational Day (G) 12. A likely systemic target of PAE is the stress peptide corticotrophin-releasing factor (CRF), as activation of CRF receptor 1 (CRFR1) in the medial nucleus of the central amygdala (CeM) is known to increase anxiety-like behavior in adults. To determine if CRF-CRFR1 interactions underly PAE-induced anxiety, functional changes in CRF system activity were investigated in adolescent male and female Sprague Dawley rats following G12 PAE. Compared to air-exposed controls, PAE increased basal spontaneous (s) inhibitory post-synaptic current (IPSC) frequency in the CeM of males, but not females. Furthermore, PAE blunted CRFR1-regulated miniature (m) IPSCs in a sex- and dose-specific manner, and only PAE males demonstrated tonic CRFR1 activity in the CeM. It was further determined that G12 PAE decreased CRFR1 mRNA in the CeM of males while increasing regional expression in females. Finally, infusion of a CRFR1 agonist into the CeM of adolescents produced a blunted expression of CRFR1-induced anxiety-like behavior exclusively in PAE males, mirroring the blunted physiology demonstrated by PAE males. Cumulatively, these data suggest that CRFR1 function within the CeM is age- and sex-specific, and PAE not only increases the expression of anxiety-like behavior, but may reduce the efficacy of treatment for PAE-induced anxiety through CRFR1-associated mechanisms. Therefore, future research will be necessary to develop targeted treatment of anxiety disorders in individuals with a history of PAE.

## Introduction

Children exposed to alcohol *in-utero* can suffer from numerous physical, cognitive and behavioral impairments classified under the term Fetal Alcohol Spectrum Disorders (FASD). FASDs are estimated to affect 1-5% of the U.S. population [1, 2], rates which are likely conservative due to underreporting of maternal drinking habits in response to societal stigma [3]. In children with confirmed cases of prenatal alcohol exposure (PAE), 21% meet clinical diagnostic criteria for anxiety disorders [4], with symptomology evident from infancy throughout adolescence [5]. This increased anxiety-like behavior has been recapitulated by multiple PAE paradigms in animal models, which vary in both severity of alcohol exposure and timing of exposure during gestation (see review: [6]). We have shown that a single exposure to a moderate dose of ethanol on gestational day (G)12 of pregnancy in rats increased generalized anxiety-like behaviors in adolescent male, but not female, offspring [7]. Although PAE-induced behavioral deficits, such as anxiety, have been well established across the lifespan [4], the neurobiological mechanisms by which PAE increases generalized anxiety is yet unknown.

Importantly, the amygdala, a brain structure associated with regulation of affect, including anxiety-like behaviors [8], undergoes significant development from G10-13 in rodent models [9]. Within the amygdalar complex, the medial central amygdala (CeM) serves as the output center for the amygdala, projecting to multiple downstream brain regions responsible for expression of anxiety-like behaviors [10]. Perturbations to the CeM under alcohol-naïve conditions have repeatedly altered expression of anxiety-like behaviors in rodent and primate models [11-13]. As G12 PAE increases anxiety-like behavior in adolescent offspring, the CeM is a likely target of PAE-induced impairments that underlie increased anxiety-like behavior.

Importantly, this region is rich in concentrations of corticotrophin-releasing factor (CRF) and its receptor, CRFR1 [14-16], which are established targets of PAE [17-19]. CRFR1 activity regulates local GABAergic neurotransmission, predominantly by increasing GABA release via presynaptic mechanisms in ethanol-naïve adult males [20]. Interestingly, we recently discovered this activity is both age and sex-specific, with adolescent males and females exhibiting *decreased* GABA release following CRFR1 activation [21]. Additionally, males with a history of adult alcohol exposure demonstrate increased sensitivity to CRF and CRFR1 antagonists [20]. Importantly, PAE throughout the entire gestational period produces subnucleus-specific, age- and sex-dependent alterations to CRF and CRFR1 expression throughout the amygdala [19, 22, 23]. However, it is unknown if PAE produces *functional* changes to the CRF system in the CeM, and if these changes may specifically promote the expression of FASD-characteristic anxiety-like behavior.

Therefore, the objective of the present study was to investigate whether G12 PAE would produce changes in CRFR1 function and expression in the CeM, leading to altered expression of anxiety-like behavior in exposed adolescents. We discovered that moderate PAE produces sex-specific changes in basal GABAergic activity and CRFR1 function. We further determined that PAE is associated with sex-specific expression of CRFR1 mRNA within the CeM, as assessed by RNAscope *in-situ* hybridization. Finally, using site-specific pharmacological manipulations, we found that infusion of a CRFR1 agonist into the CeA altered anxiety-like behavior in a sex- and exposure-dependent manner in adolescents, including a PAE-like increase in anxiety-like behavior in ethanol-naïve males.

## Methods

For the expanded and detailed methodology, please refer to *Supplementary Materials*.

### Animals and G12 PAE Paradigm

Adult male and female Sprague Dawley breeders were obtained from Envigo/Harlan (Indianapolis, IN) and permitted to acclimate at least one week prior to breeding. On G12, dams were placed in vapor inhalation chambers and were exposed to either room air (control group) or a vaporized ethanol (experimental group) as previously shown [7]. Exposures lasted for 6 h total (9:00-15:00), at which point dams were returned to colony to carry out their pregnancies. Pups were weaned on P21 and housed with same-sex littermates until experimentation. To avoid carryover effects, subjects were randomly assigned to only one experimental investigation (whole cell electrophysiology, RNAscope *in-situ hybridization* [*ISH*], or behavioral pharmacology assessment). All animal procedures were approved by the Binghamton University Institutional Animal Care and Use Committee.

### Whole-cell Electrophysiology

All slice electrophysiology experiments - including slice preparation, whole-cell current-clamp recordings, and whole-cell voltage-clamp recordings – were conducted in P40-48 offspring and mirror recently-published experimental procedures investigating CRFR1-regulated GABAergic activity within the CeM of alcohol-naïve adolescent Sprague Dawley rats [21].

### RNAscope in-situ hybridization (ISH)

Tissue slices containing the CeM were collected from G12-exposed adolescent male and female rats (P40-48) and stained with riboprobes targeting 1) CRFR1 mRNA (ACD: NM_030999.3) and 2) two cell-type biomarkers: Somatostatin (SST) (ACD: NM_001048.3) and Calbindin (CB) (ACD: NM_031984.2). Slides were also counterstained with nuclear DNA-labelling DAPI (GenBank: EF191515.1). SST and CB were selected as cell markers because of previously established localization within the CeM [24-27], and co-localization with CRFR1 [28, 29]. Stained tissue was scanned into digital images for quantitative HALO imaging analysis platform (Indica Labs) of riboprobe+ cells.

### Intracerebral cannulations and CRFR1 agonist infusion

G12-exposed offspring (P36-39) received bilateral guide cannula implantations into the CeA, recovering for 5 days prior to behavioral testing. On the day of testing, cannulated rats of each exposure/sex were randomly assigned to receive either ACSF (vehicle group) or 100 nM CRFR1 agonist Stressin-1 (experimental group). Subjects were infused bilaterally with a volume of 0.25 μl per side over 1 min using Hamilton glass syringes and an infusion pump (Harvard Apparatus, Holliston, MA). Rats were permitted 10 min rest prior to light-dark box (LDB) testing.

### Light-Dark Box (LDB)

As a validated assessment of anxiety-like behaviors in rodents [30, 31], our lab has previously used the LDB to measure behavioral changes in adolescents following G12 PAE [7]. This procedure was again performed in G12 subjects, now following 100 nM Stressin-1 or ACSF infusion. Video-recorded activity (15 min) between light and dark chambers was later assessed for group differences in behavior.

### Experimental Design and Statistical Analysis

To avoid experimenter bias, all data analyses were conducted by an individual blind to the conditions of the subject. All statistical analyses, including *t*-tests and between-subject analyses of variance (ANOVA), were performed using GraphPad 8 Software (Prism), with significance defined as *p* ≤ 0.05. Full statistics for ANOVAs, including *F*-values, *p*-values, degrees of freedom and group sample sizes, are detailed in supplementary materials. In the event of significant main effects or interactions, *post hoc* Sidak’s multiple comparison tests were performed to determine specific group differences. All data are presented as mean ± standard error of the mean (SEM).

## Results

Extended and detailed results, including all statistical information, are available in the supplemental materials.

### Membrane properties of CeM neurons are sex and exposure-specific

Assessment of membrane properties revealed a significant effect of exposure on neuronal membrane resistance, which was independent of sex (*Supplementary Table 1*). In both sexes, PAE neurons exhibited lower resistances than air-exposed controls (*p*=0.036). Although there was no effect of exposure on membrane capacitance, there was a main effect of sex (*p*=0.049), with neurons in females exhibiting higher capacitances than males.

### Both sex and prenatal exposure influence the excitability of CeM neurons

In a current-neutral configuration, cells were assessed for resting membrane potential (RMP). Sex did not influence RMP, however a main effect of exposure (*p*=0.045) revealed a PAE-induced depolarization of RMP (*Supplementary Table 1*). Cells were then held at -70 mV to normalize and assess differences in excitability across groups. From cells that were responsive to current injection, there were no differences in rheobase across exposures and sexes. However, exposure did significantly influence the membrane potential at first AP (*p*=0.035), with PAE groups demonstrating lower thresholds than controls, an effect driven primarily by PAE males (*Supplementary Table 1*). This did not correspond with a significant main effect of sex, or an exposure x sex interaction.

Quantification of firing activity revealed a significant exposure x sex interaction (*p=*0.013), whereupon PAE males exhibited significantly reduced activity compared to control males (*Fig. 1A,C*), with no significant effect of exposure in females (*Fig. 1B,C*). We also found a significant main effect of exposure when examining time to first AP (*p*=0.007), with PAE groups responding quicker to current injection than control groups (*Supplementary Table 1*), regardless of sex. AP amplitude statistically differed between sexes (*p*<0.001), with females demonstrating greater amplitudes than males, independent of exposure. Females also demonstrated longer AP half-widths than males in recorded cells independent of exposure.

**Figure 1.**
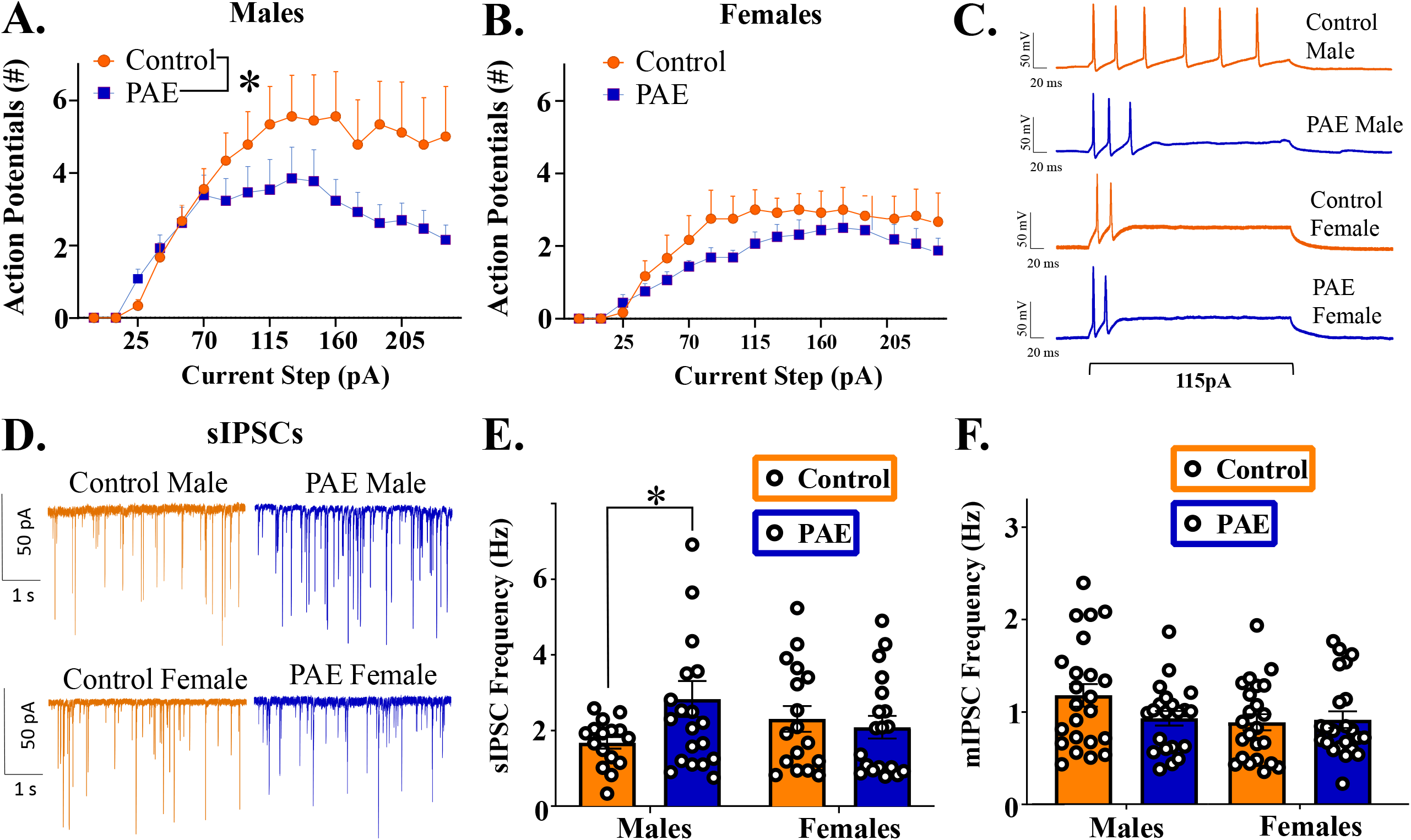
Excitability and native IPSCS of neurons within the CeM across exposure and sex. Increasing current steps (pA) produced significantly more firing activity in control males than PAE males (A,C), with no difference in females between control and PAE groups (B,C). (D) Representative sIPSC activity of neurons in the CeM: there was a significant effect of exposure only in males (E), with PAE males exhibiting significantly greater sIPSC frequency than air-exposed males. There were no effects of exposure or sex on mIPSC frequency (F). * indicates significant difference between groups (*p* < 0.05)

### Basal synaptic transmission in the CeM is sex- and exposure-specific

To determine if exposure and sex influenced basal inhibitory synaptic activity in the CeM, both spontaneous and miniature inhibitory postsynaptic currents (sIPSCs and mIPSCs) were recorded. As represented in *Fig. 1D-E*, analyses of basal sIPSC frequency revealed a significant interaction of exposure x sex (*p*=0.040). Post-hoc analyses revealed significantly higher sIPSC frequency in PAE males compared to sex-matched controls (*p=*0.039), whereas exposure produced no differences in sIPSC frequency in females. In contrast, action potential-independent mIPSC frequency was not impacted by exposure or sex (*Fig. 1F)*. Assessment of sIPSC and mIPSC amplitude revealed no significant effects of sex, nor a significant sex x exposure interaction in CeM neurons (*Supplementary Figure 1*).

### PAE blunts CRFR1 modulated GABAergic activity in a sex- and dose-dependent manner

Given our previous findings of sex-specific CRFR1-regulated mIPSC activity within the CeM of naïve adolescents [21], CRFR1-regulated activity was analyzed independently in each sex. To determine if regulation of GABA transmission by CRFR1 is altered by PAE, we assessed the effect of Stressin-1 (10 nM, 100 nM and 1 μM) on mIPSCs. Analyses of drug effects were analyzed by % change in activity from baseline, as detailed below. Changes in raw frequency values were also statistically assessed, and significant raw value comparisons mirrored significant % changes from baseline (*Supplementary Table 2*).

In males, analysis of Stressin-1-induced changes in mIPSC frequency revealed a significant main effect of dose of drug (*p*<0.001), as well as an interaction between prenatal exposure and dose of drug (*p*=0.018; *Fig. 2A,B*). Post-hoc analyses revealed no significant difference between control and PAE males at the 10 nM dose, and 10 nM Stressin-1 did not produce a significant change in mIPSC frequency in either group. At 100 nM however, drug effects were significantly different between exposure groups; specifically, a significant reduction in mIPSC frequency in control subjects (*p*<0.001) was prominently blunted in PAE subjects (*p*=0.019; *Fig. 2C*). However, this exposure effect was no longer present at the 1 μM concentration, as there were significant and comparable reductions in mIPSC frequency in both control males (*p*=0.002) and PAE males (*p*<0.001) at this dose. In females, analysis of Stressin-1-induced changes in mIPSC frequency revealed a significant main effect of dose of drug (*p*=0.019), as well as a main effect of exposure (*p*=0.018; *Fig. 2D)*. Post-hoc analyses determined that exposure-specific effects were dose-dependent (*Fig. 2E*). At 10 nM, there was no significant effect of drug in either control or PAE groups. 100 nM significantly attenuated exhibited a of mIPSC frequency to the same extent in both control (*p*=0.012) and PAE (*p*=0.005) groups. However, while 1 μM significantly reduced mIPSC frequency in control females (*p*=0.005), this effect was absent in PAE females (*Fig. 2F*), resulting in a significant effect of exposure at this dose (*p=*0.004).

**Figure 2.**
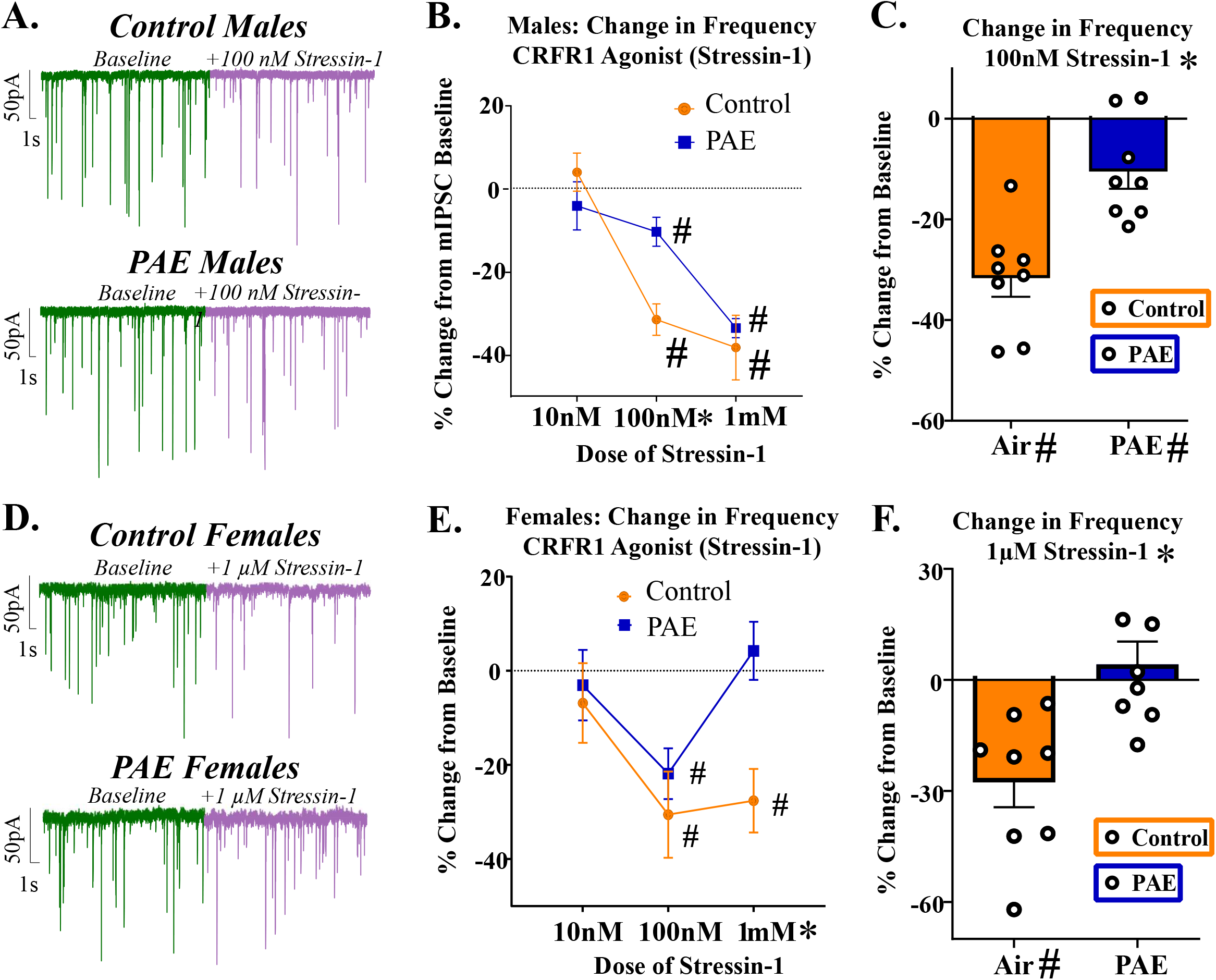
Change in mIPSCs following bath application of selective CRFR1 agonist, Stressin-1, in males (A-C) and females (D-F). (A-B) CRFR1-regulated changes in mIPSC frequency in adolescent males, reported as % change from baseline activity. CRFR1 activation produces significant dose-dependent changes in both control and PAE adolescents. In control animals, 100nM and 1µM Stressin-1 significantly decreased mIPSC frequency to similar degrees, while PAE animals exhibit a blunted decrease at 100nM which is recovered at the highest dose. (C) Bar graph of % change in mIPSC frequency from control and PAE males following bath application of 100nM Stressin-1, which revealed a significant effect of exposure in a blunted attenuation of mIPSC frequency in PAE males. (D-E) CRFR1-regulated changes in mIPSC frequency in females, reported as % change from baseline activity. 100nM Stressin-1 significantly attenuated mIPSC frequency in both control and PAE females. However, 1µM Stressin-1 resulted in a loss of that attenuation in PAE females, while control females continue to show that consistent reduction in mIPSC frequency. (F) Bar graph and representative traces of % change in mIPSC frequency from control and PAE females following bath application of 1µM Stressin-1. * indicates significant effect of exposure (*p* < 0.05) # signifies significant difference from 0.

In both males and females, analyses of % change in mIPSC amplitude, as well as raw value changes, revealed no significant effects of exposure or dose of drug (*Supplementary Fig. 1*).

### PAE increases tonic CRFR1 activity exclusively in males

To assess tonic activation of CRFR1 in the CeM, the CRFR1-selective antagonist, NBI 35965 (1 μM) was bath applied following baseline recordings of mIPSCs. Analysis of change in mIPSC frequency revealed a significant effect of exposure in males (*p*<0.001), whereupon NBI significantly potentiated mIPSC frequency in PAE males (*p*=0.001) without producing a change in control males (*Fig. 3A)*. In females, exposure did not change response to NBI, and neither control nor PAE females exhibited a significant change in mIPSC frequency from baseline (*Fig. 3B)*. Analysis of change in mIPSC amplitude in these same cells revealed no significant main effects of exposure in males or females in response to NBI (*Supplementary Fig. 1*).

**Figure 3.**
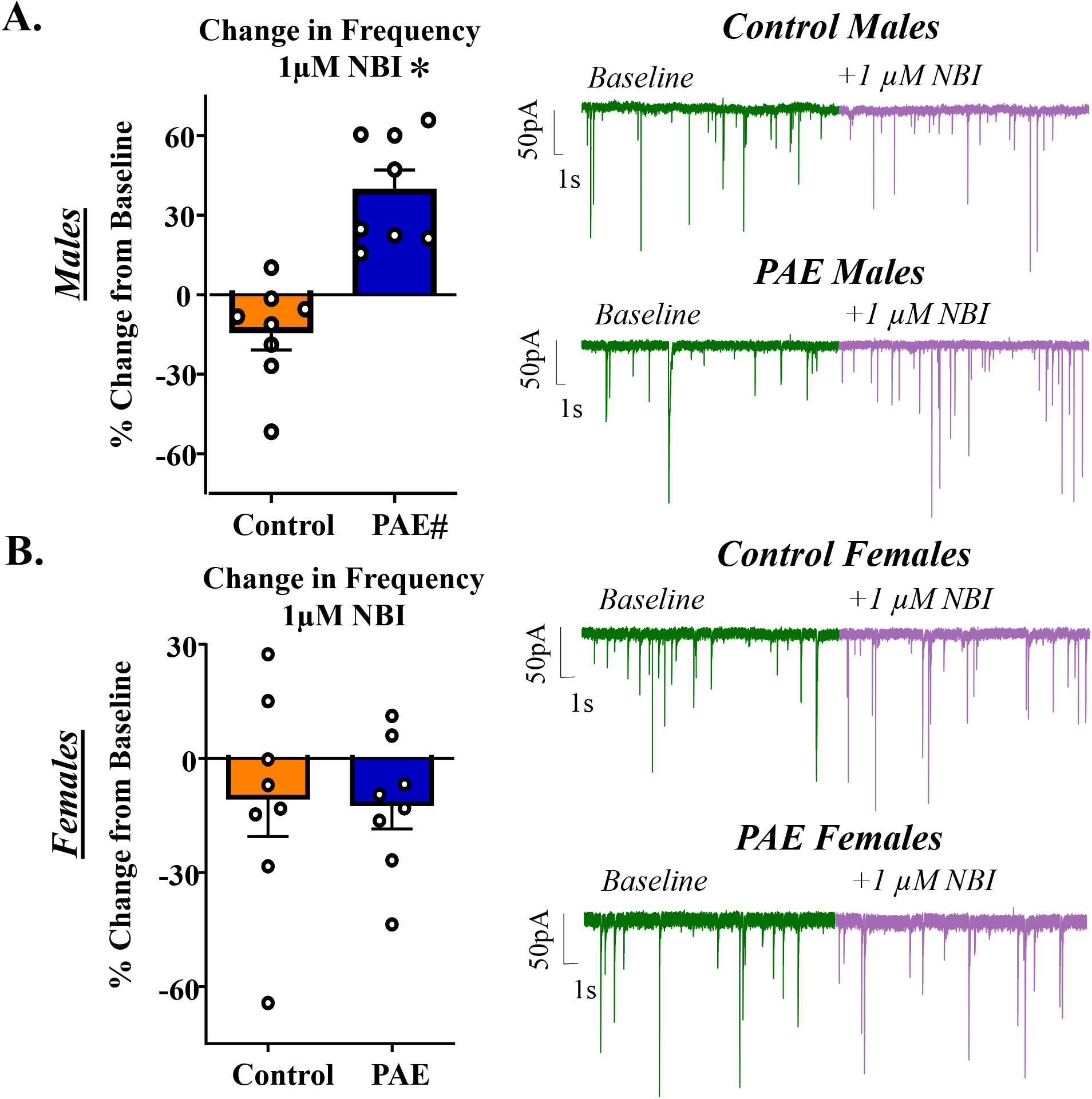
Males and females: change in mIPSCs following bath application of selective CRFR1-receptor antagonist, NBI (1µM). (A) mIPSC frequency activity before and after CRFR1 blockade in both sexes. (B) mIPSC frequency activity following CRFR1 blockade, reported as % change in baseline activity. Only PAE males demonstrate significant tonic activation of CRFR1. * indicates significant effect of exposure (*p* < 0.05) # signifies significant difference from 0.

### PAE produces sex-specific changes in the expression of CRFR1 mRNA in the CeM

We next evaluated whether G12 moderate PAE influenced the expression of CRFR1 mRNA within the CeM. We first determined that PAE did not change DAPI stained nuclei concentrations within this region across exposure or sex (*Fig*. 4A). Next, we found that quantification of raw mRNA transcript abundance mirrored the patterns of expression observed in co-localization of mRNA with DAPI-stained nuclei (*Supplementary Figure 3*); therefore, all data are subsequently presented by cell+ expression. The following comparisons were selected *a priori* for nested analyses of cellular subtypes, with *p*-values corrected for multiple comparisons: 1) Control Males vs Control Females, 2) Control Males vs PAE Males and 3) Control Females vs PAE females.

**Figure 4.**
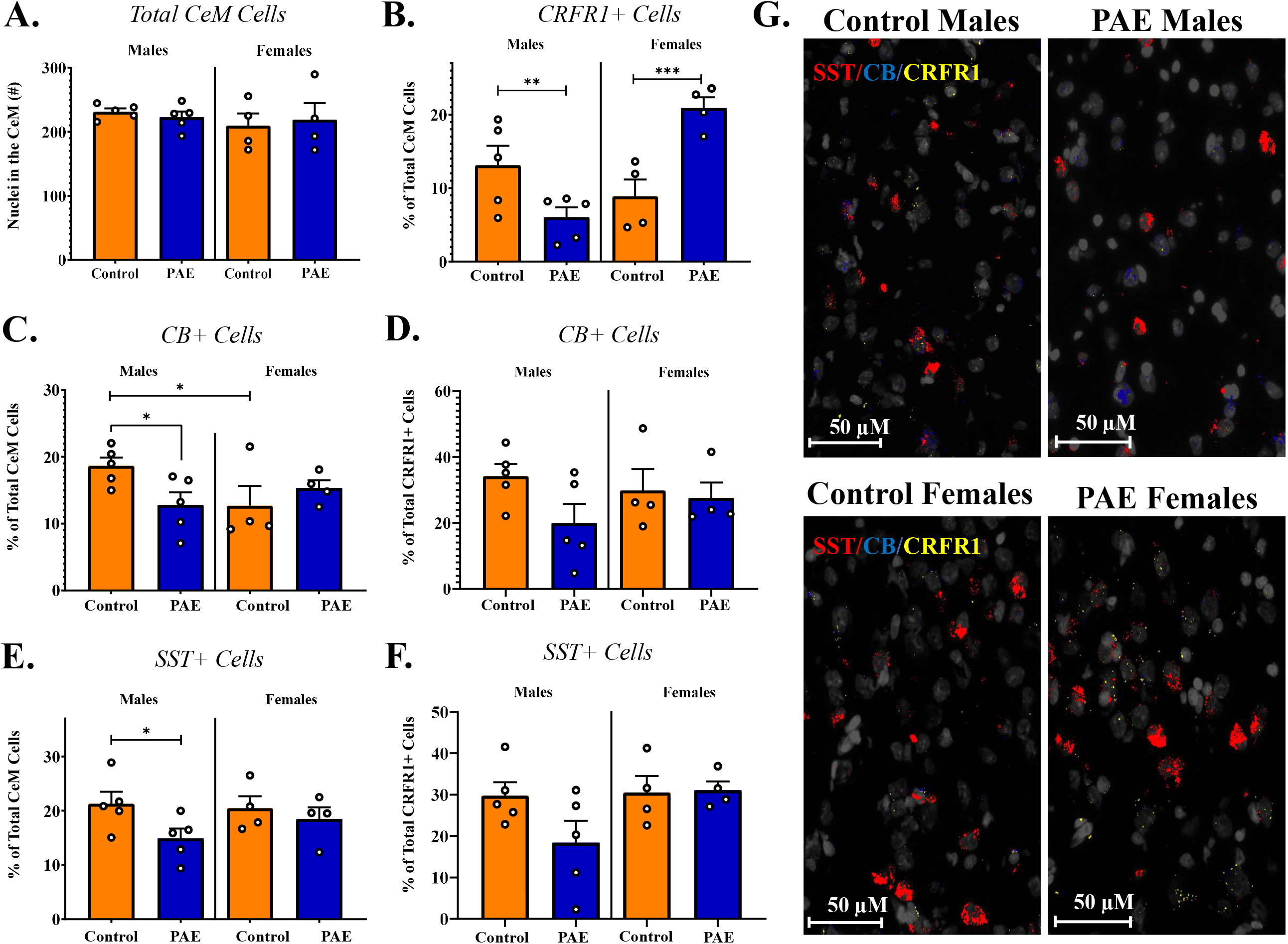
mRNA quantification in the CeM of prenatally exposed male and female adolescents. (A) Quantification of nuclei within this region. Neither sex nor exposure changed the # of nuclei counterstained by DAPI. (B) Quantification of CRFR1+ cells, reported as % of nuclei-stained cells. PAE significantly decreased the # of CRFR1+ cells in the CeM of males, while significantly increasing CRFR1+ cells in females. (C) Quantification of CB+ cells, reported as % of nuclei-stained cells. In controls, males exhibited higher proportions of CB+ cells than females. PAE significantly decreased the proportion of CB+ cells in the CeM in males, without affecting females. (D) Quantification CB+ cells co-labelled with CRFR1 mRNA. Factors of sex and exposure do not change the proportion of CRFR1/CB+ cells in the CeM. (E) Quantification of SST+ cells in the CeM, reported as % of nuclei-stained cells. PAE significantly decreased the proportion of SST+ cells in the CeM in males, without affecting females. (F) Quantification SST+ cells co-labelled with CRFR1 mRNA. Factors of sex and exposure do not change the proportion of CRFR1/SST+ cells in the CeM (G) Representative images (40X) of fluorescently labelled mRNA in CeM cells across sex and exposure. Red: SST. Blue: CB. Yellow: CRFR1.. * indicates significant effect of exposure (*p* < 0.05), ** (*p < 0*.*01*), *** (*p* < 0.001)

In quantification of CRFR1+ cells, control males and females did not differ in their proportional expression in CeM cells (*Fig. 4B)*. However, exposure produced significant changes in % CRFR1+ cells within this region in both males (*p*=0.001) and females (*p*<0.001). Importantly, this change was bidirectional between sexes, with PAE producing a significant reduction in CRFR1+ cells in males, and a significant increase in CRFR1+ cells in females.

Quantification of total CB+ cells within this region revealed a significant effect of sex in control animals, with control females exhibiting lower proportions of CB+ cells than males (*Fig. 4C)*. PAE did not change the proportion of CB+ cells within this region in females, however PAE significantly reduced the percentage of CB+ cells in males (*p*=0.017). In CRFR1+ cells, approximately 1/3 of cells co-labeled with CB in both male and female controls, with no statistical difference between the two groups. PAE did not significantly change the proportion of CRFR1+ cells co-labeled with CB in females or males, although males demonstrated a non-significant trend toward a PAE-induced decrease in CRFR1-CB co-labelled cells (*p*=0.079; *Fig. 4D)*.

Quantification of total SST+ cells within this region revealed no difference in proportion of SST+ cells between control males and females (*Fig. 4E)*. PAE did not change the proportion of SST+ cells within this region in females, however PAE significantly reduced the percentage of SST+ cells in males (*p*=0.031). In CRFR1+ cells, approximately 30% of cells co-labeled with SST in both male and female controls, with no statistical difference between the two groups. PAE did not significantly change the proportion of CRFR1+ cells co-labeled with SST in females or males (*Fig. 4F)*. There was minimal overlap between SST+ and CB+ cells within this region (<5% across experimental groups), and therefore double-labeled SST/CB cells were not statistically analyzed due to insufficient power.

### PAE produces sex-specific changes in behavioral response to CRFR1 agonist infusion into the CeM

In our final series of experiments, we evaluated whether moderate PAE changed behavioral response to the activation of CRFR1 within the CeM via a region-targeted infusion of 100 nM Stressin-1. This concentration was selected because it produced exposure-specific physiological response in males, but not females, in our electrophysiology assessments of mIPSC frequency (*Fig. 2)*. We used the LDB assay to assess multiple measures of generalized anxiety-like behavior.

When assessing time spent in the light chamber of the apparatus, a significant interaction between drug infusion (ACSF, Stressin-1) and exposure was uncovered in males (*p*=0.022; *Fig. 5A)*, an interaction that was not statistically significant in females. To further elaborate on sex-specific effects, males were separated graphically by exposure to depict group-specific drug effects (*Figs. 5B,C,G,H)* and females were separated graphically by drug infusion to depict group-specific exposure effects (*Figs. 5D,E,I,J)*. Follow-up analyses of time spent in the light chamber revealed that in males, Stressin-1 infusion in control animals reduced overall time spent in the light chamber (*p*=0.014; *Fig. 5A&B)* but did not change time spent in the light chamber in PAE males (*Fig. 5A&C)*. Control males and PAE males did not statistically differ in their time spent in light chamber when infused with ACSF, although there is a notable reduction in time spent in the light chamber by PAE males (*p*=0.094). In females, there was no effect of Stressin-1 infusion in either control or PAE females; however, PAE females infused with ACSF demonstrated significantly less time in the light chamber than control females infused with ACSF (*p*=0.025; *Fig. 5A&D)*. This difference was not present between exposure groups following infusion of Stressin-1 (*Fig. 5A&E)*.

**Figure 5A-E.**
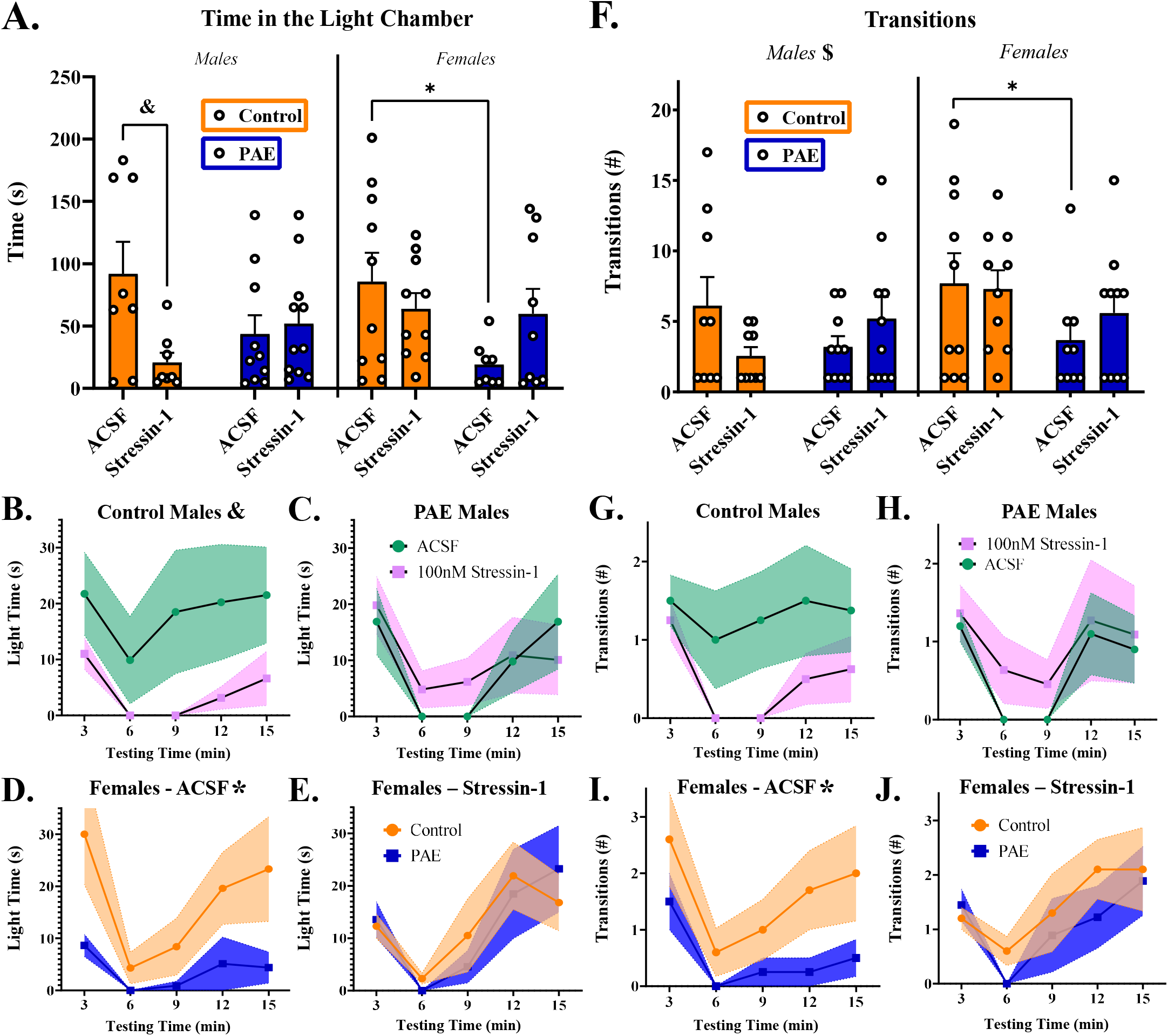
Light time (s) in the LDB between exposures, sexes and infusions of either ACSF or 100nM Stressin-1 into the CeA. (A) Total time spent in the light chamber of the LDB assay. There was a significant drug effect in control males, with Stressin-1 infusion reducing the time spent in the light chamber. In females, there was a significant effect of exposure only in ACSF-infused animals, with PAE reducing time spent in the light chamber. (B) Time spent in the light chamber by control males, across 3min time bins of the test. Stressin-1 infused males spend significantly less time in the light chamber than ACSF-infused males. (C) Time spent in the light chamber by PAE males, across 3min time bins of the test. Drug infusion did not change time spent in the light in PAE males. (D) Time spent in the light chamber by females infused with ACSF, across 3min time bins of the test. PAE females spent significantly less time in the light chamber than control females. (E) Time spent in the light chamber by females infused with Stressin-1, across 3min time bins of the test. There was no effect of exposure on light time in Stressin-1-infused females. **Figure 5F-J.** Transitions between light and dark chambers in the LDB. (F) Males demonstrated a significant exposure x drug infusion interaction, producing a Stressin-1-induced reduction in transitions in control males and a Stressin-1-induced increase in transitions in PAE males. (G) Transitions in the LDB by control males, across 3min time bins of the test. ACSF-infused males demonstrate a consistent rate of transitions throughout the test, whereas Stressin-1-infused males demonstrate fewer, less consistent transitions. (H) Transitions in the LDB by PAE males, across 3min time bins of the test. Drug infusion did not change transition patterns in PAE males. (I) Transitions in the LDB in ACSF-infused females, across 3min time bins of the test. Control females transition more throughout the test than PAE females. (J) Transitions in the LDB in Stressin-1-infused females, across 3min time bins of the test. Exposure did not change transition patterns in response to Stressin-1 infusion. & indicates significant effect of Stressin-1 (*p* < 0.05), * indicates significant effect of exposure, $ indicates a significant interaction between drug infusion and prenatal exposure

When assessing the number of transitions between light and dark chambers, males demonstrated a significant exposure x drug infusion interaction (*p=*0.048; *Fig. 5F)*, but this interaction was not significant in females. There were no main effects of drug infusion in either sex, and no main effect of exposure in males. Although not statistically significant, there was a notable trend toward a main effect of exposure in females driven by PAE (*p*=0.084), with PAE reducing the number of transitions throughout the test (*Fig. 5F&I)*. There was no significant effect of exposure in females infused with Stressin-1 (*Fig. 5F&J*). Post-hoc analyses in males revealed a non-significant reduction in transitions following infusion of Stressin-1 in control males (*p* = 0.075; *Fig. 5F&G)* with no effect in PAE males (*Fig. 5F&H)*.

When examining transitions across time bins, we observed that control males infused with ACSF demonstrated consistent rates of crossing between chambers throughout the testing period. In contrast, control males infused with Stressin-1 did not transition for at least 9 min after initially entering the dark chamber. To determine if transitioning behavior was time-specific, we ran additional post-hoc analyses investigating changes in transition rates between time bins. There was no significant effect of time in control males infused with ACSF, but there was a significant effect of time in control males infused with Stressin-1 (*p*=0.037; *Fig. 5G)* that was absent in PAE males.

When assessing time to first enter the dark chamber (egress latency), there were no significant effects of drug infusion or exposure in males (*Supplementary Fig. 5A)*. In females, there was also no effect of drug infusion, however, there was a significant effect of exposure. Post-hoc analyses determined this significant effect of exposure was once again specific to ACSF-infused females (*p*=0.018), with PAE reducing egress latency in this group, while exposure did not change response to Stressin-1 infusion in females. Assessments of time to first return to the dark chamber (re-entry latency) revealed no significant effects of drug infusion or exposure in males or females (*Supplementary Fig. 5B)*. Assessment of head poking from the dark chamber into the light chamber revealed similar null results in both sexes (*Supplementary Fig. 5C)*.

## Discussion

In the present investigation, PAE reduced CeM neuron firing and potentiated sIPSC frequency in male offspring with no effect of exposure on mIPSCs. Changes in sIPSCs, but not mIPSCs, point to altered mechanisms upstream of the presynaptic terminal. The CeA contains both CRFR1+ and CRFR1-neurons, which display distinct tonic activity mediated by different GABA-A receptor subunits [15]. Importantly, acute ethanol enhances δ subunit-mediated sIPSCs in CRFR1-neurons, suggesting that shifts in cell type/expression may contribute to alcohol-induced sIPSC potentiation. Notably, as G12 PAE reduced overall CRFR1+ cells in the CeM in our *ISH* assessment, it is possible that observed sIPSC frequency enhancement is attributable to increased proportions of CRFR1-neurons in this region. Additionally, sIPSC effects could be attributed to greater cumulative input onto recorded CeM neurons from local or distal GABAergic projections. Since we found that PAE males exhibited reduced firing activity through both stimulated and under current-neutral conditions (data not shown), the source of increased sIPSCs is likely outside of the CeM.

As previously observed in drug-naïve adolescent males [21], a moderate (100 nM) concentration of a CRFR1-selective agonist significantly attenuated mIPSC frequency (but not amplitude) in control males, an effect which was not further amplified at a higher (1 µM) concentration. Importantly, the attenuation of mIPSC frequency in PAE males at this moderate concentration was blunted ∼66% compared to controls, but fully recovered at the highest dose. If CeM CRFR1 receptor activity contributes to stress-responses and anxiety-like behavior in adolescent males, as attributed to adult males [32, 33], this rightward-shift in response suggests more CRFR1 activation may be required to produce an appropriate stress-response in PAE males. Changes in mIPSC frequency, but not amplitude, indicate altered function at the presynaptic terminal; therefore, we hypothesized that PAE-blunted activity in males may be attributable to reduced presynaptic CRFR1 expression. Consistent with our hypothesis, we found reduced CRFR1 mRNA in PAE males compared to controls, an effect which was not specific to CB+ or SST+ cells. Interestingly, our data also revealed a PAE-induced reduction in CB+ and SST+ cells within the CeM of males, without changing overall nucleic concentrations. These findings suggest that PAE alters additional sex-specific, CRFR1-independent mechanisms in the CeM without depreciating overall cell quantities.

In contrast, PAE females demonstrated an attenuated response to the CRFR1 agonist at the highest concentration (1µM), suggesting that regulation by CRFR1 receptors was either inactivated or compensated for at this highest concentration. Importantly, PAE also increased the expression of CRFR1 mRNA within the CeM in this same group. Across sexes, PAE produced opposing directional shifts in CRFR1 mRNA that do not correspond with opposing directional changes in mIPSC frequency – rather, both PAE males and females demonstrate attenuated mIPSC frequency, albeit at different concentrations of CRFR1 agonist. Together, these data suggest that, although PAE may alter CRFR1 mRNA expression, this alone is not sufficient to explain the sex-specific modulation of GABAergic activity uncovered in our study, and future studies should further examine the locus of these functional alterations.

Interestingly, across sexes and exposures, only PAE males demonstrated significant tonic CRFR1-regulated activity in the CeM. Bath application of the CRFR1 antagonist significantly potentiated mIPSC frequency in PAE males, opposite the attenuation observed with the selective agonist, which may indicate higher levels of endogenous CRF. Increased expression of CRF mRNA has been repeatedly reported in preclinical alcohol exposure models, including acute [34, 35], chronic [36, 37] and prenatal exposures [17, 23]. In drug-naïve adult rats, a recent histological study has reported sex and region-specific expression of extrahypothalamic CRF protein, mirroring sex-specific findings in CRF mRNA investigations [38]. Future research should specifically determine whether G12 PAE increases endogenous CRF within the CeM.

Given that our electrophysiology data revealed sex and dose-specific response to moderate levels of CRFR1 agonist (100 nM Stressin-1), we predicted that microinjection of Stressin-1 at this concentration would produce distinct anxiety-like behavior between control and PAE males, but not between control and PAE females. Consistent with our hypothesis, control males demonstrated significantly increased anxiety-like behavior following Stressin-1 infusion, spending less time in the light chamber and transitioning less frequently/consistently between chambers than subjects infused with ACSF. This Stressin-1-induced increase in anxiety-like behavior was absent in PAE males, that demonstrated comparable performance across LDB measures regardless of drug infusion (ACSF or Stressin-1). This significant interaction between PAE and Stressin-1-infusion in males was absent in females, as PAE did not produce behavioral changes in response to Stressin-1. Although not significant, PAE males injected with ACSF spent less time in the light side relative to control males injected with ACSF, suggesting a basal anxiety-like phenotype, potentially due to tonic activation of CeM CRFR1 that is consistent with the findings from our NBI experiment. Interestingly, these findings in males replicate our previous LDB investigation using this model of PAE in non-surgerized adolescent offspring [7]. Although adolescent females did not demonstrate a PAE-induced change in behavioral response to Stressin-1, PAE females infused with ACSF exhibited increased anxiety-like behavior, suggesting that females may be impacted by prenatal exposure through non-CRFR1-regulated mechanisms.

## Conclusion

G12 PAE produces neurophysiological and behavioral impairments in exposed adolescent offspring, with both sexes demonstrating unique susceptibility to changes in CRFR1 function. These sex-specific vulnerabilities may be attributable to alcohol’s established effects on GABAergic activity within the CeM, as well as differences in CRF/CRFR1 expression and associated signaling pathways. Importantly, while our findings indicate that the CRF/CRFR1 system is a target of PAE, canonical approaches of targeting CRFR1 to treat stress and anxiety disorders [39] may not translate to PAE-associated anxiety or anxiety-related disorders in females due to differences in neuroadaptations of the CRFR1 system. Thus, we must continue to investigate and develop methods for diagnosing PAE in exposed individuals, facilitating the prescription of appropriate, targeted treatment for FASD-associated anxiety disorders.

## Acknowledgements

The authors wish to acknowledge and extend their sincerest thanks to Drs. Molly Deak and Terrance Deak for their instruction and insight on *RNAscope* experiments. The experiments included in this manuscript were supported by NIAAA grants P50 AA01782306, T32 AA025606 and F31 AA028166.

## Author Contributions

**Siara Rouzer**: Conceptualization, methodology, formal analysis, investigation, writing-original draft preparation, writing-reviewing and editing, visualization, funding acquisition.

**Marvin Diaz**: Conceptualization, methodology, writing-reviewing and editing, supervision, project administration, funding acquisition.

## Figure Legends

**Supplementary Table 1**. Cell properties of CeM neurons across sex and prenatal exposure, reported as Mean (SEM). PAE neurons demonstrated higher membrane resistance compared to control animals, independent of sex, while females demonstrated greater membrane capacitance than males, independent of exposure. Compared to control groups, PAE groups demonstrated more depolarized resting membrane potentials, lower action potential thresholds and quicker time to production of the 1^st^ AP following current injection. The amount of current required to produce this first AP, the rheobase, did not differ between sex or exposure groups. AP amplitudes were higher in females than males, and AP half-widths were shorter in males than females, and both of these effects were unaffected by prenatal exposure. & indicates significant main effect of exposure (*p* < 0.05), # indicates significant main effect of sex (*p* < 0.05).

**Supplementary Table 2**. Paired t-test comparisons of frequency values (Hz) at baseline (mIPSC) and following drug application in all experimental groups.

**Supplementary Figure 1**. mIPSC amplitudes across all CRFR1-targetting experiments/groups. (A) Amplitude values in males across exposure and drug/dose. There were no significant effects of exposure or drug/dose on mIPSC amplitude. (B) Amplitude values in females across exposure and drug/dose. There were no significant effects of exposure or drug/dose on mIPSC amplitude. (C) % Change in mIPSC amplitude per experimental group for each CRFR1-targeted experiment. There were no significant effects of exposure on mIPSC amplitude at any dose/drug in either sex.

**Supplementary Figure 2**. Representative full-sized images (40X) of fluorescently labelled mRNA in CeM cells of males and females across exposure. Red: SST. Blue: CB. Yellow: CRFR1.

**Supplementary Figure 3**. Quantification of total transcript copies in the CeM of prenatally exposed male and female adolescents. (A) Quantification of CRFR1 transcript within the CeM. PAE significantly decreased CRFR1 transcript levels in the CeM of males, while significantly increasing CRFR1 transcript levels in females. (B) Quantification of CB transcript within the CeM. In controls, males exhibited higher levels of CB transcript than females. Furthermore, PAE significantly decreased CB transcript concentrations in males, without affecting females. (C) Quantification of SST transcript in the CeM. PAE significantly decreased the levels of SST transcript in males, without affecting females. * indicates significant effect of exposure (*p* < 0.05), ** (*p < 0*.*01*), *** (*p* < 0.001), **** (*p* < 0.0001)

**Supplementary Figure 4**. Guide cannula targeting the CeA were bilaterally implanted during surgery in G12 adolescents. On the day of testing in the light-dark box, implanted-subjects were infused with either ACSF (control) or 100nM Stressin-1.

**Supplementary Figure 5**. Behavioral measures in the LDB between exposures, sexes and infusions of either ACSF or 100nM Stressin-1 into the CeA. A) Latency to leave the light chamber for the first time (Egress Latency). PAE shortens egress latency only in ACSF-infused females. B) Latency to re-enter the dark chamber for the first time (Re-Entry Latency). There were no effects of exposure or drug infusion on this measure in either males or females. C) Head pokes from the dark chamber into the light chamber during LDB testing. There were no effects of exposure or drug infusion on this measure in either males or females. * indicates significant effect of exposure (*p* < 0.05)

