## Supplementary Materials for "Moderate prenatal alcohol exposure modifies sex-specific CRFR1 activity in the central amygdala and anxiety-like behavior in adolescent offspring"

| | Control Males ($n$ = 9-16) | Control Females ($n$ = 12-17) | PAE Males ($n$ = 13-15) | PAE Females ($n$ = 16-19) |
|---|---|---|---|---|
| **Membrane Resistance (MΩ) &** | 387.38 (47.35) | 482.66 (45.47) | 437.21 (37.86) | 533.32 (52.22) |
| **Membrane Capacitance (pF) #** | 49.22 (3.43) | 45.45 (3.44) | 41.89 (3.05) | 39.18 (2.76) |
| **Resting Membrane Potential (mV) &** | -75.19 (2.27) | -70.34 (2.26) | -69.22 (1.59) | -67.20 (2.29) |
| **AP Threshold (mV) &** | -64.00 (1.14) | -61.90 (1.67) | -59.10 (1.33) | -60.51 (1.33) |
| **Rheobase (pA)** | 51.67 (3.33) | 48.75 (5.37) | 45.00 (2.82) | 46.56 (4.32) |
| **Time to 1st AP (ms) &** | 107.06 (13.36) | 101.83 (14.08) | 87.36 (10.89) | 61.37 (2.19) |
| **AP Amplitude (mv) #** | 82.13 (5.09) | 95.86 (2.46) | 74.22 (2.94) | 94.50 (2.92) |
| **AP Half-Width (ms) #** | 1.99 (0.14) | 2.22 (0.12) | 2.13 (0.15) | 2.55 (0.17) |

**Supplementary Table 1.** Cell properties of CeM neurons across sex and prenatal exposure, reported as Mean (SEM). PAE neurons demonstrated higher membrane resistance compared to control animals, independent of sex, while females demonstrated greater membrane capacitance than males, independent of exposure. Compared to control groups, PAE groups demonstrated more depolarized resting membrane potentials, lower action potential thresholds and quicker time to production of the 1st AP following current injection. The amount of current required to produce this first AP, the rheobase, did not differ between sex or exposure groups. AP amplitudes were higher in females than males, and AP half-widths were shorter in males than females, and both of these effects were unaffected by prenatal exposure. & indicates significant main effect of exposure ($p < 0.05$), # indicates significant main effect of sex ($p < 0.05$).

| Experimental Group | Drug | Baseline frequency | + Drug Frequency | *t-value* | *p-value* | Sig? ($p < 0.05$) |
|---|---|---|---|---|---|---|
| **Control Males** | 10nM Stressin-1 | 0.830 (0.12) | 0.827 (0.14) | 0.912 | 0.392 | No |
| | **100nM Stressin-1** | **1.200 (0.18)** | **0.800 (0.11)** | **4.328** | **0.003** | **Yes** |
| | **1μM Stressin-1** | **1.793 (0.33)** | **1.152 (0.28)** | **4.155** | **0.004** | **Yes** |
| | 1μM NBI | 1.249 (0.23) | 1.094 (0.16) | 1.838 | 0.116 | Yes |
| **PAE Males** | 10nM Stressin-1 | 0.672 (0.08) | 0.632 (0.09) | 1.136 | 0.293 | No |
| | 100nM Stressin-1 | 1.333 (0.32) | 1.17 (0.25) | 1.761 | 0.129 | No |
| | **1μM Stressin-1** | **1.400 (0.28)** | **0.918 (0.18)** | **4.423** | **0.003** | **Yes** |
| | **1μM NBI** | **0.779 (0.25)** | **0.988 (0.27)** | **6.401** | **< 0.001** | **Yes** |
| **Control Females** | 10nM Stressin-1 | 0.809 (0.20) | 0.670 (0.12) | 1.511 | 0.174 | No |
| | **100nM Stressin-1** | **1.146 (0.08)** | **0.770 (0.10)** | **2.902** | **0.023** | **Yes** |
| | **1μM Stressin-1** | **0.650 (0.10)** | **0.436 (0.04)** | **2.409** | **0.047** | **Yes** |
| | 1μM NBI | 0.778 (0.24) | 0.593 (0.14) | 1.082 | 0.315 | No |
| **PAE Females** | 10nM Stressin-1 | 1.210 (0.32) | 1.150 (0.29) | 0.856 | 0.420 | No |
| | **100nM Stressin-1** | **0.928 (0.14)** | **0.686 (0.07)** | **2.561** | **0.038** | **Yes** |
| | 1μM Stressin-1 | 0.908 (0.16) | 0.985 (0.15) | 1.403 | 0.203 | No |
| | 1μM NBI | 1.088 (0.21) | 0.901 (0.17) | 2.032 | 0.082 | No |

**Supplementary Table 2.** Paired t-test comparisons of frequency values (Hz) at baseline (mIPSC) and following drug application in all experimental groups.

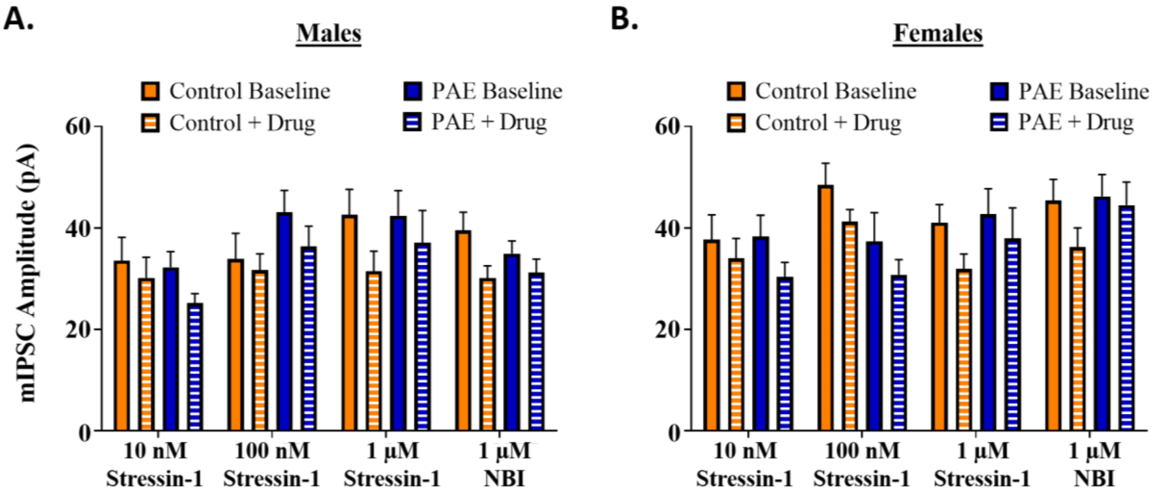


C.

| AMPLITUDE | 10nM Stressin | 100nM Stressin | 1µM Stressin | 1µM NBI |
|---|---|---|---|---|
| **Control Males (*n* = 7-8)** | -9.17 (5.47) | -11.88 (8.78) | -23.33 (8.04) | -21.55 (4.30) |
| **Control Females (*n* = 8-9)** | -7.88 (3.67) | -12.70 (4.75) | -19.90 (7.62) | -20.59 (3.43) |
| **PAE Males (*n* = 8)** | -18.65 (7.62) | -12.56 (8.22) | -12.61 (10.74) | -10.56 (4.243) |
| **PAE Females (*n* = 8)** | -19.99 (1.79) | -13.40 (5.07) | -12.88 (6.84) | -10.47 (4.01) |

**Supplementary Figure 1.** mIPSC amplitudes across all CRFR1-targetting experiments/groups. **A)** Amplitude values in males across exposure and drug/dose. There were no significant effects of exposure or drug/dose on mIPSC amplitude. **B)** Amplitude values in females across exposure and drug/dose. There were no significant effects of exposure or drug/dose on mIPSC amplitude. **C)** % Change in mIPSC amplitude per experimental group for each CRFR1-targeted experiment. There were no significant effects of exposure on mIPSC amplitude at any dose/drug in either sex.

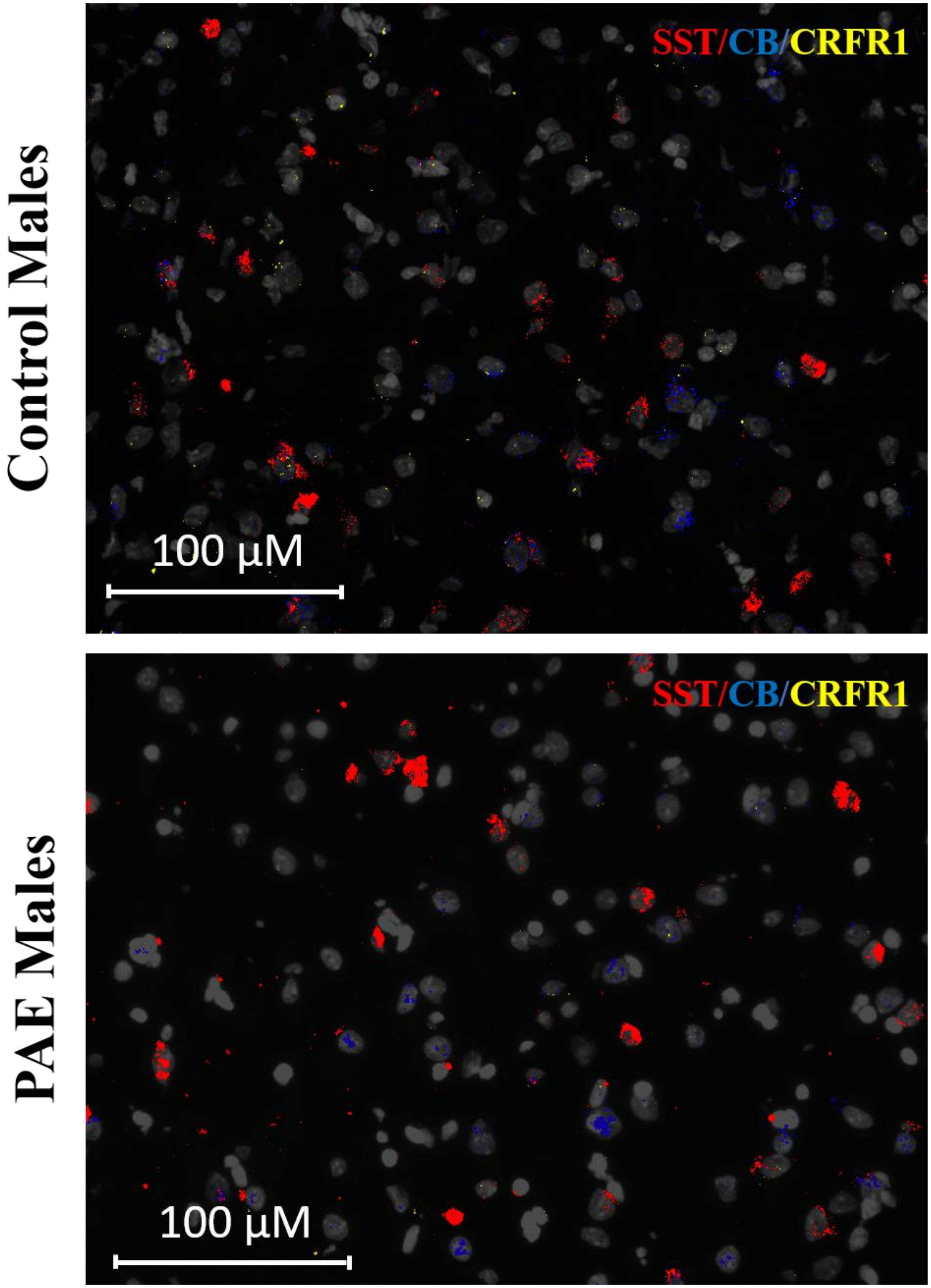


**Supplementary Figure 2a.** Representative full-sized images (40X) of fluorescently labelled mRNA in CeM cells of males across exposure. Red: SST. Blue: CB. Yellow: CRFR1.

Control Females

PAE Females

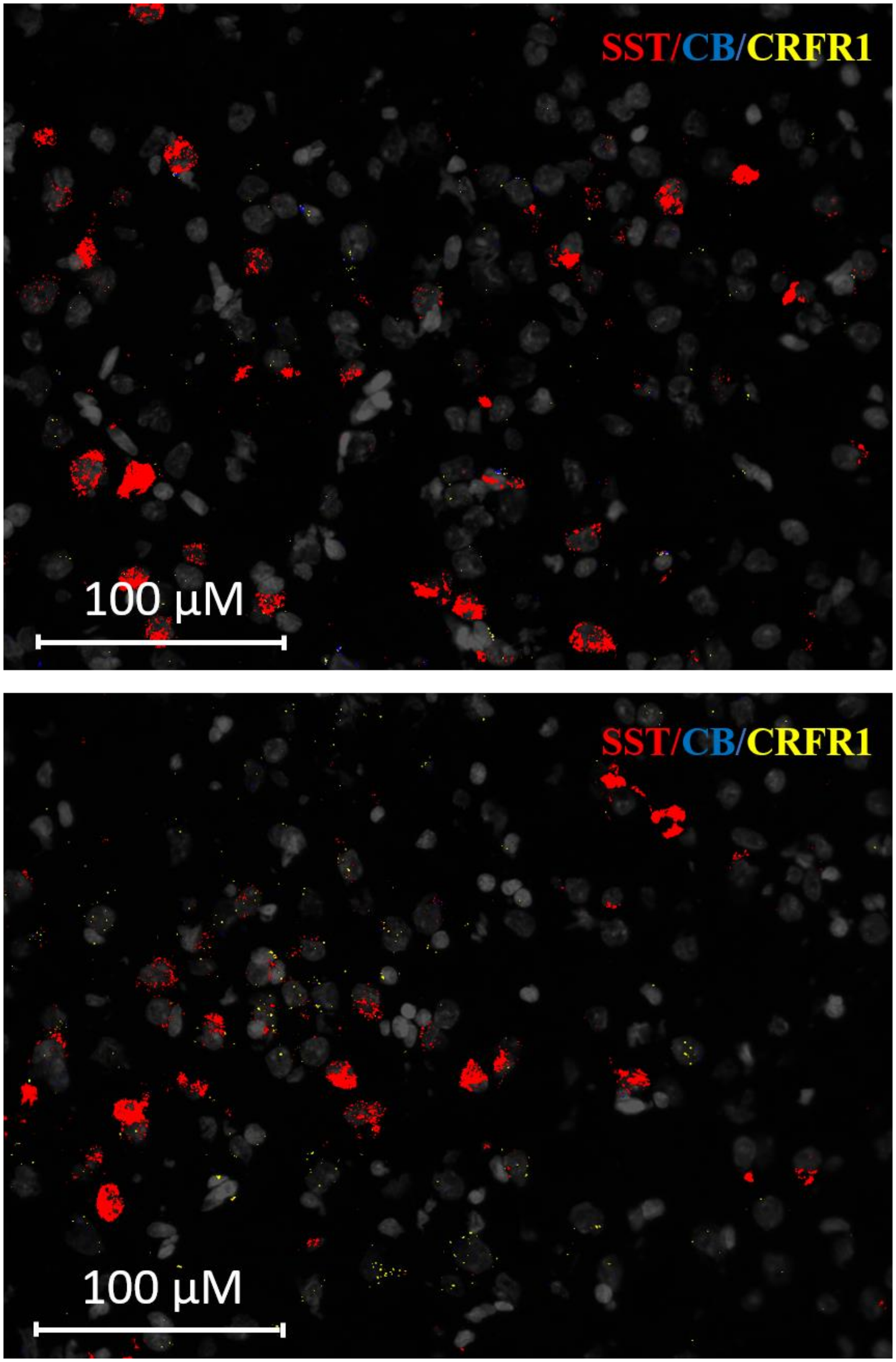


**Supplementary Figure 2b.** Representative full-sized images (40X) of fluorescently labelled mRNA in CeM cells of females across exposure. Red: SST. Blue: CB. Yellow: CRFR1

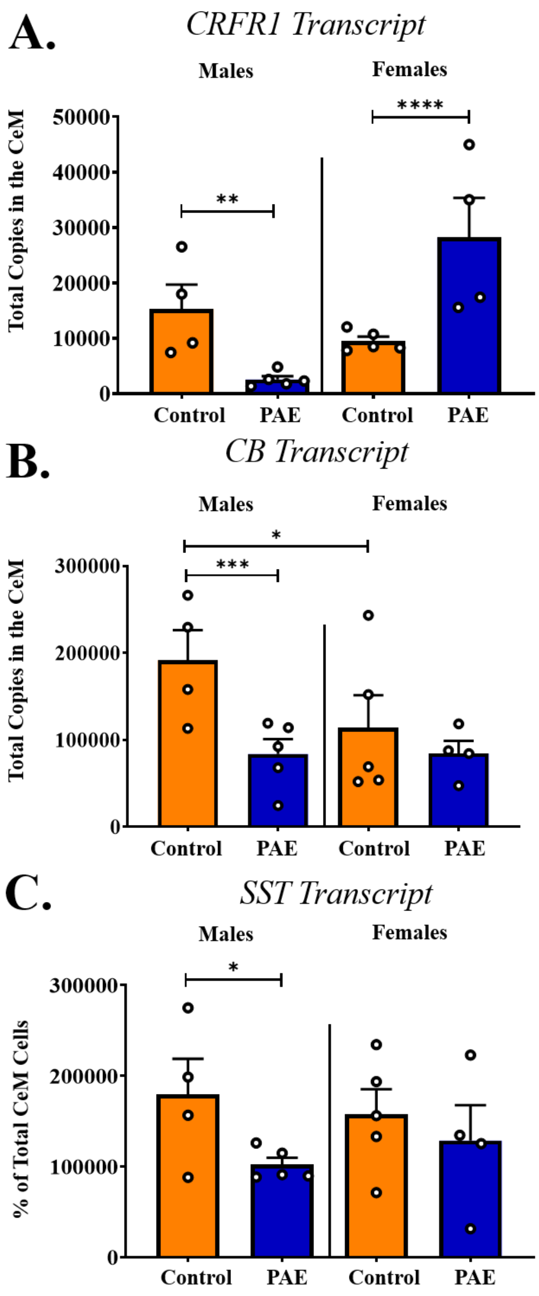


**Supplementary Figure 3.** Quantification of total transcript copies in the CeM of prenatally exposed male and female adolescents. A) Quantification of CRFR1 transcript within the CeM. PAE significantly decreased CRFR1 transcript levels in the CeM of males, while significantly increasing CRFR1 transcript levels in females. B) Quantification of CB transcript within the CeM. In controls, males exhibited higher levels of CB transcript than females. Furthermore, PAE significantly decreased CB transcript concentrations in males, without affecting females. C) Quantification of SST transcript in the CeM. PAE significantly decreased the levels of SST transcript in males, without affecting females. * indicates significant effect of exposure ($p < 0.05$), ** ($p < 0.01$), *** ($p < 0.001$), **** ($p < 0.0001$)

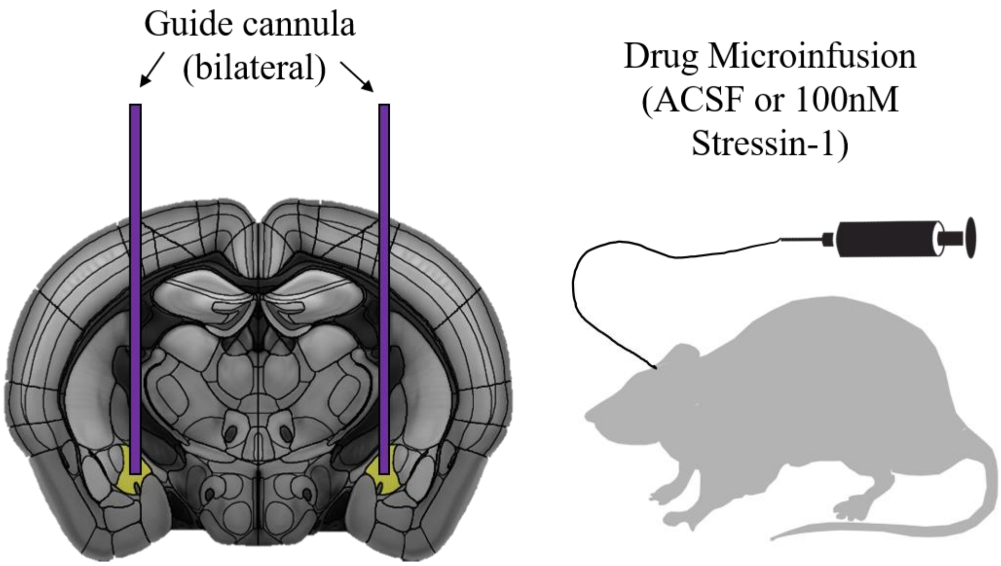


**Supplementary Figure 4.** Guide cannula targeting the CeA were bilaterally implanted during surgery in G12 adolescents. On the day of testing in the light-dark box, implanted-subjects were infused with either ACSF (control) or 100nM Stressin-1.

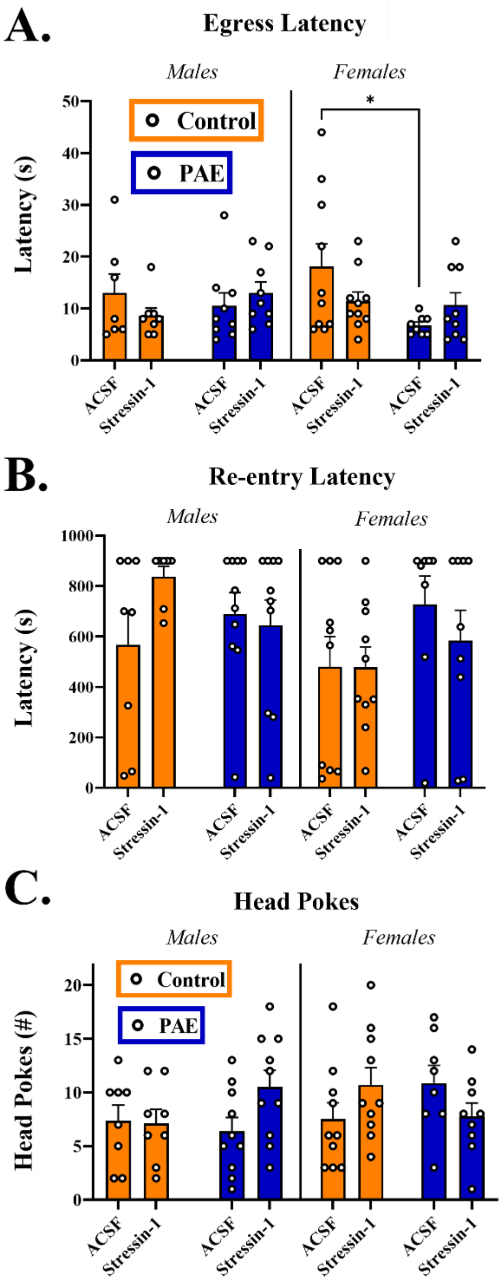


**Supplementary Figure 5.** Behavioral measures in the LDB between exposures, sexes and infusions of either ACSF or 100nM Stressin-1 into the CeA. A) Latency to leave the light chamber for the first time (Egress Latency). PAE shortens egress latency only in ACSF-infused females. B) Latency to re-enter the dark chamber for the first time (Re-Entry Latency). There were no effects of exposure or drug infusion on this measure in either males or females. C) Head pokes from the dark chamber into the light chamber during LDB testing. There were no effects of exposure or drug infusion on this measure in either males or females. * indicates significant effect of exposure ($p < 0.05$)

*Expanded Methods*

***Animals.*** To avoid introducing shipping stress during development, all experimental subjects were bred in-house as previously described [1]. Adult male and female Sprague Dawley breeders were obtained from Envigo/Harlan (Indianapolis, IN) and permitted to acclimate at least one week prior to breeding. Upon detection of sperm in vaginal smears (gestational day (G) 1), pregnant dams were isolated and housed with a plastic hut and crinkle paper as nesting material. On G12, dams were placed in vapor inhalation chambers to create prenatal exposure groups, as described below. Otherwise, dams remained unhandled until delivery. After parturition, litters were culled to 5:5 males/females on Postnatal Day (P)2 and housing conditions maintained, with a plastic hut and crinkle paper. Pups were weaned from their mother on P21 and housed with same-sex littermates until experimentation. All animals were group-housed (2-3 animals per cage) in a temperature-controlled (22°C) vivarium and maintained on a 12:12 h light:dark cycle (lights on at 07:00 h). Subjects were provided *ad libitum* access to food (5L0D PicoLab Laboratory Rodent diet) and water throughout the duration of experimentation. To avoid carryover effects, subjects were randomly assigned to only one experimental investigation (whole cell electrophysiology, RNAscope *in-situ hybridization* [*ISH*], or behavioral pharmacology assessment). All animal procedures were approved by the Binghamton University Institutional Animal Care and Use Committee.

***G12 Vapor Ethanol Exposure.*** As previously established in this model of prenatal exposure [1], pregnant dams in their home cages were transferred to vaporized inhalation chambers on G12 and assigned to one of two conditions: a chamber with room air (control group) or a chamber with vaporized ethanol (experimental group). Dams were not handled at any point during transfer to and from the chambers. Exposures lasted for 6 h (9:00-15:00), during which dams had free access to food and water. At 15:00, cages were removed and returned to the colony room for the remainder of pregnancy. Food from ethanol-exposed cages was replaced to avoid additional exposure to ethanol through absorption. To avoid the stress of tail-blood collection on experimental animals, a different subset of pregnant dams was sampled following G12 exposure (15:00 mark) to determine average blood ethanol concentrations; these dams were promptly euthanized after tail-blood collection. To ensure this G12 model produced reliable BECs over the course of PAE experiments, periodic assessments of ethanol levels were performed using tail-blood samples collected from an experimentally-naïve subset of pregnant dams. Furthermore, within exposure, ethanol vapor readings were recorded hourly to ensure consistency between exposures.

***Whole-cell Electrophysiology.*** Drugs and Chemicals. All chemicals and kynurenic acid were purchased from Sigma-Aldrich (St. Louis, MO, USA). APV, tetrodotoxin (TTX), Stressin-1, CGP 55845 and NBI 35965 were purchased from Tocris/R&D systems (Bristol, UK).

Slice Preparation. Adolescent (P40-48) male and female rats were sedated with a 250 mg/kg dose of ketamine and quickly decapitated, as previously performed by our lab [2]. This specific period of mid-late adolescence has been repeatedly used by our lab to investigate impairments engendered by prenatal alcohol exposure [1, 3]. Furthermore, ethanol-naïve offspring of this age range have recently demonstrated sex-specific CRFR1-regulation of IPSC activity within the CeM [4]. Brains were rapidly removed and immersed in ice-cold oxygenated (95% $O_2$-5% $CO_2$) sucrose artificial cerebrospinal fluid (ACSF) containing (in mM): sucrose (220), KCl (2), $NaH_2PO_4$ (1.25), $NaHCO_3$ (26), glucose (10), $MgSO_4$ (12), $CaCl_2$ (0.2), and ketamine (0.43). Coronal slices (300 μM) containing the CeM were collected with a Vibratome (Leica Microsystems. Bannocknurn, IL, USA). Slices were incubated in 34°C normal ACSF (in mM): NaCl (125), KCl (2), $NaH_2PO_4$ (1.3), $NaHCO_3$ (26), glucose (10), $CaCl_2$ (0.1), $MgSO_4$ (0.1), ascorbic acid (0.04), and continuously bubbled at 95% $O_2$-5% $CO_2$ for at least 40 minutes before recording. All experiments were performed within 6 hours of slice preparation.

Whole-cell patch-clamp recordings. Following incubation, slices were transferred to a recording chamber, where oxygenated ACSF was warmed to 34°C and continuously superfused over the submerged slice at 3.3 ml/min. Recording electrodes of 3-5 MΩ tip resistance were pulled from borosilicate glass capillary tubing (Sutter Instruments) using a Flaming-Brown puller (Sutter Instruments). Recordings were collected from the CeM with patch pipettes filled with experiment-specific internal solutions (see below). Membrane properties (averaged across entire recordings) were provided by the membrane test in pClamp 10 (Molecular Devices) and electrophysiology data were acquired with a MultiClamp 700B (Molecular Devices, Sunnyvale, CA) at 10 kHz, filtered at 1 kHz, and stored for later analysis using pClamp 10 software (Molecular Devices).

*Voltage-clamp recordings:* For voltage-clamp experiments, a KCl-based internal solution was used for detecting $GABA_A$ receptor-mediated currents, containing (in mM): KCl (135), HEPES (10), $MgCl_2$ (2), EGTA (0.5), Mg-ATP (5), Na-GTP (1), and QX314-Cl (1); 300 mOsm; 7.3 pH with KOH. For recordings of $GABA_A$ receptor-mediated spontaneous inhibitory postsynaptic currents (sIPSCs), AMPA and NMDA glutamate receptors were pharmacologically blocked using 1 mM kynurenic acid and 50 μM APV, respectively, and $GABA_B$ receptors were blocked with CGP 36645 (1 μM). For data collection of miniature IPSCs (mIPSCs), recordings were made in the presence of the Na+ channel blocker, TTX (1 μM). Baseline recordings of sIPSCs and mIPSCs were allowed to equilibrate for at least 5 min before recording began. A 3-min baseline period was recorded prior to drug application.

For electrophysiological assessment of CRFR1 function, the CRFR1 selective-agonist, Stressin-1 (10 nM, 100 nM and 1 μM) or the selective antagonist, NBI 35965 (1 μM), was applied for 15 min. As GABAergic neurons within the CeM exhibited generally stable access resistance, this length of time ensured that changes in transmission could be visualized for stability. To determine the timing of a stable drug effect, a time course of drug exposure was constructed during the pilot phase of experimentation and stable activity was observed 5 min following drug application. Therefore, the first 5 min of drug application was removed from final analyses. Additionally, for all experiments, only recordings with an access resistance change of <20% were included in final analyses.

*Current-clamp recordings*. For assessments of intrinsic excitability and resting membrane potential (RMP), recordings were collected from the CeM with patch pipettes filled with a K-gluconate-based internal solution containing (in mM): K-Gluconate (120), KCl (15), EGTA (0.1), HEPES (10), $MgCl_2$ (4.0), MgATP (4.0), $Na_3GTP$ (0.3) and Phosphocreatine (7.0); 280-290 mOsm; 7.25 pH with KOH. Cells were opened in a voltage-clamp configuration (holding potential = -70 mV) and switched to current-clamp settings for at least 7 minutes to allow neurons to dialyze prior to applying a series of depolarizing current steps (increments of 15 pA, 500 ms duration). Subsequent firing activity was recorded for future analysis. Rheobase, time to first action potential, and action potential threshold, peak and half-width were determined with the first action potential fired with the lowest stimulation current as assessed by pClamp 10 software (Molecular Devices). A liquid junction potential of -19.4 mV was also calculated using this software; all reported membrane potentials have been adjusted to account for this disparity.

***RNAscope in-situ hybridization (ISH).*** Adolescent male and female rats (P40-48) were euthanized with Fatal-Plus solution (100mg/kg; manufacturer: Vortech; contains pentobarbital sodium 390mg/mL, propylene glycol 0.01mg/mL, ethyl alcohol 0.29mg/mL, benzyl alcohol 0.2 mg/mL) and transcardially perfused first with phosphate buffered saline (PBS) and then 4%

paraformaldehyde (PFA) to remove all blood from the brain. Whole brains were extracted and preserved in PFA for 24 h, then transferred to a solution of 30% sucrose in PBS until sectioned.

Subsequent experimental procedures followed the manufacturer's instructions. Immediately prior to sectioning, whole brains were flash frozen and tissue sections (12 μm thick) collected and mounted on positively charged microscopic glass slides. For each slide (1 slide = 1 animal), two adjacent tissue sections were stained with riboprobes targeting 1) CRFR1 mRNA (ACD: NM_030999.3) and 2) two cell-type biomarkers: Somatostatin (SST) (ACD: NM_001048.3) and Calbindin (CB) (ACD: NM_031984.2). Slides were also counterstained with nuclear DNA-labelling DAPI (GenBank: EF191515.1). SST and CB were selected as cell markers because they can be found within the CeM [5-8], and co-localize with CRFR1 [9, 10]. Stained slides were cover-slipped with ProLong Gold Antifade Mountant (Thermo Fisher Scientific, Waltham, MA) and scanned into digital images for analysis.

RNAscope *ISH* fluorescent images of the CeM in each hemisphere were captured at 40× magnification using a BX-X800 fluorescent microscope (Keyence, Osaka, Japan). Images were saved as 16-bit TIFF files and uploaded to the HALO imaging analysis platform (Indica Labs) to perform quantitative analysis. Quantitative analysis parameters were selected as recommended by ACD, with settings optimized to detect round nuclei and discrete fluorescent dots. CRFR1+ cell concentrations are moderate within the CeM, and to avoid false-positives, detection parameters for CRFR1+ cells required multiple (3+) distinct dots for CRFR1 transcript; however, in post-analyses review of our data, all CRFR1+ cells contained at least 8 distinct dots. Once parameters of analysis were established, all images (1 image per hemisphere, 4 total images per slide) were batch-analyzed using identical settings.

***Intracerebral cannulations and CRFR1 agonist infusion.*** On P36-39, both air and ethanol-exposed offspring were anesthetized with isoflurane (as inhalant: 3% induction, 2% maintenance) and stereotaxically implanted bilaterally with 23-gauge stainless steel guide cannulas (Plastics One, Roanoke, VA) targeting the CeA in adolescent rats: −2.2 mm posterior to bregma, +/− 4.2 mm from the midline and −6.3 mm ventral from the surface of the skull (*Supplementary Fig. 4*). Guide cannulas were secured to the skull with dental cement and anchor screws, after which subjects recovered for 5 days prior to behavioral testing. For 48 h postoperatively, surgerized subjects received intraperitoneal injections of the analgesic buprenorphine (0.03mg/kg) every 12 h. Animals were also weighed and handled daily postoperatively to assess their health and acclimate them to handling by the experimenter prior to behavioral testing. On the day of behavioral testing, cannulated rats of each exposure/sex were randomly assigned to receive either ACSF (vehicle group) or 100 nM CRFR1 agonist Stressin-1 (experimental group). Subjects were infused bilaterally with a volume of 0.25 μl per side over 1 min using Hamilton glass syringes and an infusion pump (Harvard Apparatus, Holliston, MA). Injectors were removed from guide cannulas 1 min after the end of the infusions, and rats were permitted 10 min rest prior to light-dark box (LDB) testing.

***Light-Dark Box (LDB).*** As a validated assessment of anxiety-like behaviors in rodents [11, 12], our lab has previously used the LDB to measure behavioral changes in adolescents following G12 PAE [1]. This procedure was again performed in G12 subjects, now following Stressin-1 infusion. Briefly, subjects were placed into the "light box" of the dual-chambered apparatus, facing away from the open aperture bridging this box with the "dark box". The researcher quickly exited the room, and subjects were given 15 min to freely explore the apparatus. This behavior was video-recorded for later analyses of 1) time spent in each chamber, 2) the latency to first enter the dark chamber (egress latency), 3) latency to return to the light box for the first time (re-entry latency), 4) the number of “head pokes” into the aperture while staying within the dark box and finally 5) the number of transitions between chambers.

Immediately following LDB testing, subjects were euthanized with intraperitoneal injections of Fatal Plus (100 mg/kg). Indigo dye was infused through guide cannula prior to brain extraction. Brains were then frozen for later visual verification of cannula placement on a cryostat (CM1510; Leica Biosystems). Only subjects who demonstrated successful bilateral targeting of the CeA were included in final experimental analyses.

***Experimental Design and Statistical Analysis.*** Based on preliminary data, a power analysis (*G*Power 3.1.9.2*) for a two-way (sex x age) analysis of variance (ANOVA) and an alpha of 0.05 suggested a sample size of 8 units per group. To minimize litter effects in our sampling population, no more than 2 subjects per litter were used for any experimental group for all procedures. To avoid experimenter bias, all data analyses were conducted by an individual blind to the conditions of the subject. All statistical analyses were performed using GraphPad 8 Software (Prism) and in all assessments, significance was defined as $p \leq 0.05$. All data were assessed for outliers using the ROUT method of regression (GraphPad 8 Software, Prism), with an FDR of 1%, and all identified outliers were removed from statistical analyses. All data are presented as mean ± standard error of the mean (SEM) unless otherwise specified. In the event of significant main effects or interactions, *post hoc* Sidak’s multiple comparison tests were performed to determine specific group differences.

In electrophysiology experiments, the unit of analyses was cell, with no more than 2 cells per animal used for any given experiment. pClamp (company, City, State) and MiniAnalysis (company, City, State) were used to analyze electrophysiology recordings. Measures of membrane properties, RMP, intrinsic excitability, and basal s/mIPSCs were analyzed with 2 (sex: male, female) x 2 (prenatal exposure: air, alcohol) between-subjects ANOVA. Concentration-response activity following CRFR1 activation were reported as % change from baseline mIPSC activity and analyzed independently in each sex using 2 (prenatal exposure) x 3 (concentration: 10 nM, 100 nM, 1 μM) between-subjects ANOVAs. To determine if a drug effect was statistically significant, we used a one sample *t-test* to compare % change in activity to a null change (0). In within-cell analyses between baseline mIPSCs and drug application, effect of drug is reported as significant only when a statistical comparison between baseline activity and drug application differs by $p < 0.05$.

Statistical assessment of mRNA expression was performed using a Nested ANOVA. For each animal, two bi-hemispheric brain slices containing the CeM provided 4 total data points for each subject. These data points were then nested to account for within-subject variability and compiled into experimental groups for statistical comparison ($n$ = 4-5 animals per sex/exposure). In assessments of the LDB, we made an *a priori* decision to analyze sexes separately, as our previous behavioral investigation using the same PAE model uncovered sex-specific behavior in the LDB, which we wanted to target separately. All measures of the LDB test were analyzed using a 2 (exposure: control or PAE) x 2 (drug infused: ACSF or 100nM Stressin-1) between-subjects ANOVA.

*Detailed Results*

***Membrane properties of CeM neurons are sex and exposure-specific.*** GABAergic neurons in the CeM were identified by their approximate membrane resistance to membrane capacitance values, as previously reported [13]. The average access resistance of electrophysiological recordings was 17.63 (± 0.593), with no difference in access resistance between experimental groups. Assessment of membrane properties revealed a significant effect of exposure on neuronal membrane resistance [$F(1,63) = 4.593$, $p = 0.036$, $n$ = 15-19 cells per group], which was independent of sex [sex: $F(1,63) = 1.052$, $p = 0.309$; exposure x sex interaction: $F(1,63) = 0.025$, $p = 0.875$, $n$ = 15-19 cells per group] (*Supplementary Table 1*). In both sexes, PAE neurons exhibited lower resistances than air-exposed controls. Although there was no effect of exposure on membrane capacitance [$F(1,63) = 1.112$, $p = 0.296$, $n$ = 15-19 cells per group], there was a main effect of sex [$F(1,63) = 4.035$, $p = 0.049$, $n$ = 15-19 cells per group] and no interaction between the two variables [$F(1,63) < 0.001$, $p = 0.993$, $n$ = 15-19 cells per group]. Independent of exposure, neurons of females exhibited higher capacitances than males.

***Both sex and prenatal exposure influence the excitability of CeM neurons.*** In a current-neutral configuration, cells were assessed for resting membrane potential (RMP). Sex did not influence resting membrane potential [sex: $F(1,46) = 2.397$, $p = 0.128$; exposure x sex interaction: $F(1,46) = 0.406$, $p = 0.527$, $n$ = 9-16 cells per group], however a main effect of exposure revealed a PAE-induced depolarization of RMP [$F(1,46) = 4.229$, $p = 0.045$, $n$ = 9-16 cells per group] (*Supplementary Table 1*). Cells were then held at -70 mV to assess and normalize differences in excitability across groups. From cells that were responsive to current injection, there were no differences in rheobase across exposures and sexes [exposure: $F(1,46) = 1.022$, $p = 0.317$; sex: $F(1,46) = 0.024$, $p = 0.878$; exposure x sex interaction: $F(1,46) = 0.262$, $p = 0.612$] (*Supplementary*

*Table 1*). However, exposure did significantly influence the membrane potential at first AP [$F(1,46) = 4.707$, $p = 0.035$], with PAE groups demonstrating lower thresholds than controls, an effect driven primarily by PAE males (*Supplementary Table 1*). This did not correspond with a significant main effect of sex [$F(1,46) = 0.056$, $p = 0.813$], or an exposure x sex interaction [$F(1,46) = 1.466$, $p = 0.232$].

Quantification of firing activity revealed a significant exposure x sex interaction [$F(1,782) = 6.220$, $p = 0.013$], whereupon PAE males exhibited significantly reduced activity compared to control males [$t(1,25) = 2.188$, $p = 0.038$] (*Fig. 1A,C*). This effect of exposure was absent in females [$t(1,32) = 1.624$, $p = 0.114$] (*Fig. 1B,C*). We also found a significant main effect of exposure when examining time to first AP, with PAE groups responding quicker to current injection than control groups [$F(1,42) = 8.047$, $p = 0.007$] (*Supplementary Table 1*). This activity was not influenced by sex, either as a main effect [$F(1,42) = 2.168$, $p = 0.148$] or as an interacting variable [$F(1,42) = 0.959$, $p = 0.333$]. AP amplitude statistically differed between sexes [$F(1,46) = 26.160$, $p < 0.001$], with females demonstrating greater amplitudes than males, and no effect of exposure [$F(1,46) = 1.945$, $p = 0.170$] or an interaction of exposure x sex [$F(1,46) = 0.972$, $p = 0.329$]. Females also demonstrated longer AP half-widths than males in recorded cells [$F(1,46) = 4.126$, $p = 0.048$], independent of exposure [exposure: $F(1,46) = 2.041$, $p = 0.160$; exposure x sex interaction: $F(1,37) = 0.408$, $p = 0.526$].

***Basal synaptic transmission in the CeM is sex- and exposure-specific.*** To determine if factors of exposure and sex influenced basal inhibitory synaptic activity in the CeM, both sIPSCs and mIPSCs were recorded. As represented in *Fig. 1D-E*, analyses of basal sIPSC frequency revealed a significant interaction of exposure x sex [$F(1,67) = 4.402$, $p = 0.040$, $n = 16\text{-}19$ cells], with no main effects of exposure [$F(1,67) = 2.180$, $p = 0.145$] or sex [$F(1,67) = 0.077$, $p = 0.782$].

To account for within-animal variability of sIPSC activity across experimental groups, multiple sIPSC recordings were acquired from the same animal when possible. To avoid the inflation of statistical power from this oversampling of cells, we also collapsed cells within-animal and re-performed this analysis. The sex x PAE interaction was similarly observed when cells were grouped by animal [$F(1,27) = 5.497$, $p = 0.027$, $n$ = 6-9 animals] (data not shown). Post-hoc analyses revealed significantly higher sIPSC frequency in PAE males compared to sex-matched controls ($p = 0.039$), whereas exposure produced no differences in sIPSC frequency in females ($p = 0.884$). Assessment of sIPSC amplitude revealed no significant effects of sex [$F(1,67) = 0.772$, $p = 0.383$] or exposure [$F(1,67) = 0.956$, $p = 0.332$], nor a significant sex x exposure interaction [$F(1,67) = 0.148$, $p = 0.702$]. In contrast, action potential-independent mIPSC frequency was not impacted by exposure [$F(1,86) = 1.298$, $p = 0.258$], sex [$F(1,86) = 2.586$, $p = 0.112$], or an interaction between the two variables [$F(1,86) = 2.031$, $p = 0.158$]. Similarly, neither exposure [$F(1,86) = 0.893$, $p = 0.347$] nor sex [$F(1,86) = 0.001$, $p = 0.974$] changed mIPSC amplitude in CeM neurons, and there was no significant interaction between these variables (*Fig. 1F)* [$F(1,86) = 0.008$, $p = 0.929$].

***PAE blunts CRFR1 modulated GABAergic activity in a sex- and dose-dependent manner.*** Given our previous findings of sex-specific CRFR1-regulated mIPSC activity within the CeM of naïve adolescents [4], CRFR1-regulated activity was subsequently analyzed independently in each sex. Recently, our lab uncovered. To determine if this regulation of GABA transmission by CRFR1 is altered by PAE, we assessed the effect of Stressin-1 (10 nM, 100 nM and 1 μM) on mIPSCs. Analyses of drug effects were analyzed by % change in activity from baseline, as detailed below. Changes in raw frequency values were also statistically assessed;

significant raw value comparisons mirrored significant % changes from baseline, and have been summarized in *Supplementary Table 2*.

In males, analysis of Stressin-1-induced changes in mIPSC frequency revealed a significant main effect of dose of drug [$F(2,43) = 26.55$, $p < 0.001$], as well as an interaction between prenatal exposure and dose of drug [$F(2,43) = 4.412$, $p = 0.018$] (*Fig. 2A,B*). Post-hoc analyses revealed no significant difference between control and PAE males at the 10 nM dose [$t(1,43) = 1.158$, $p = 0.584$], whereupon 10 nM Stressin-1 did not produce a significant change in mIPSC frequency in either control [$t(1,7) = 1.015$, $p = 0.344875$, $n = 8$ cells/4 litters] or PAE adolescents [$t(1,7) = 0.825$, $p = 0.436$, $n = 8$ cells/5 litters]. At 100 nM however, drug effects were significantly different between exposure groups [$t(1,43) = 3.031$, $p = 0.012$]; specifically, the significant reduction in mIPSC frequency in control subjects [$t(1,7) = 8.419$, $p < 0.001$, $n = 8$ cells/4 litters] was prominently blunted in PAE subjects [$t(1,7) = 3.022$, $p = 0.019$, $n = 8$ cells/5 litters] (*Fig. 2C*). However, this exposure effect was no longer present at the 1 μM dose [$t(1,43) = 0.671$, $p = 0.880$], as there were significant and comparable reductions in mIPSC frequency in both control males [$t(1,7) = 4.942$, $p = 0.002$, $n = 8$ cells/4 litters] and PAE males [$t(1,7) = 14.560$, $p < 0.001$, $n = 8$ cells/4 litters] at this dose.

Analysis of drug-induced changes in mIPSC amplitude in these same cells revealed no significant main effects of exposure [$F(1,43) = 2.172$, $p = 0.148$] or dose of drug [$F(2,43) = 0.255$, $p = 0.776$] in males, nor a significant exposure x dose of drug interaction [$F(2,43) = 0.884$, $p = 0.421$] (*Supplementary Fig. 2*). Analysis of raw value changes in mIPSC amplitude revealed similar null results: mIPSC amplitudes in adolescent males were not influenced by dose of CRFR1 agonist [$F(2,42) = 2.294$, $p = 0.114$], prenatal exposure [$F(1,43) = 0.438$, $p = 0.512$] or an interaction of these two variables [$F(2,43) = 0.784$, $p = 0.467$].

In females, analysis of Stressin-1-induced changes in mIPSC frequency revealed a significant main effect of dose of drug [$F(2,42) = 4.370$, $p = 0.019$], as well as a main effect of exposure [$F(1,42) = 6.061$, $p = 0.018$] (*Fig. 2D)*. Post-hoc analyses determined that exposure-specific effects were dose-dependent (*Fig. 2E*). At 10 nM, there was no significant effect of drug in either control [$t(1,7) = 0.812$, $p = 0.444$, $n = 8$ cells/5 litters] or PAE females [$t(1,7) = 0.693$, $p = 0.693$, $n = 8$ cells/4 litters]. At 100 nM, both groups exhibit a significant attenuation of mIPSC frequency [control: $t(1,7) = 3.340$, $p = 0.012$, $n = 8$ cells/4 litters; PAE: $t(1,7) = 4.062$, $p = 0.005$, $n = 8$ cells/4 litters] that did not differ between groups [$t(1,14) = 0.819$, $p = 0.426$]. However, at 1 μM, while control females continued to exhibit a significant reduction in mIPSC frequency [$t(1,7) = 4.087$, $p = 0.005$, $n = 8$ cells/4 litters], PAE females no longer show any change from baseline frequency [$t(1,7) = 0.684$, $p = 0.5159$, $n = 8$ cells/4 litters]. The effect of exposure at this dose was statistically significant [$t(1,14) = 3.480$, $p = 0.004$] (*Fig. 2F*).

Analysis of mIPSC amplitude in these same cells revealed no significant main effects of exposure [$F(1,42) = 0.109$, $p = 0.743$] or dose of drug [$F(2,42) = 0.126$, $p = 0.882$] in females, nor a significant age x drug interaction [$F(2,42) = 1.922$, $p = 0.158$] (*Supplementary Fig. 1*). Analysis of raw value changes in mIPSC amplitude revealed similar null results: mIPSC amplitudes in adolescent males were not influenced by the dose of CRFR1 agonist [$F(2,42) = 0.645$, $p = 0.530$], prenatal exposure [$F(1,42) = 0.741$, $p = 0.394$] or an interaction of these two variables [$F(2,43) = 1.678$, $p = 0.199$].

***PAE increases tonic CRFR1 activity exclusively in males.*** To assess tonic activation of CRFR1 in the CeM, the CRFR1-selective antagonist, NBI 35965 (1 μM) was bath applied following baseline recordings of mIPSCs (*Fig. 3)*. Analysis of change in mIPSC frequency revealed a significant effect of exposure in males [$t(1,13) = 5.422$, $p < 0.001$], whereupon NBI significantly potentiated mIPSC frequency in PAE males [$t$(1,7) = 5.395, $p$ = 0.001, $n$ = 8 cells/5 litters] without producing a change in mIPSC frequency in control males [$t$(1,6) = 1.957, $p$ = 0.098, $n$ = 8 cells/5 litters]. In females, exposure did not change response to NBI [$t$(1,14) = 1.080, $p$ = 0.886], such that neither control nor PAE females exhibited a significant change in mIPSC frequency from baseline [control: $t$(1,7) = 0.1.080, $p$ = 0.316, $n$ = 8 cells/5 litters; PAE: $t$(1,7) = 2.001, $p$ = 0.086, $n$ = 8 cells/4 litters].

Analysis of change in mIPSC amplitude in these same cells revealed no significant main effects of exposure in males [$t$(1,13) = 1.815, $p$ = 0.093] or females [$t$(1,13) = 1.932, $p$ = 0.076] in response to the CRFR1 antagonist (*Supplementary Fig. 1*). Similarly, changes in raw mIPSC amplitude values did not differ by exposure in males [$F$(1,13) = 0.004, $p$ = 0.795] or females [$F$(1,14) = 0.701, $p$ = 0.416].

***PAE produces sex-specific changes in the expression of CRFR1 mRNA in the CeM.*** We next evaluated whether G12 moderate PAE influenced the expression of CRFR1 mRNA within the CeM (*Fig. 4)*. We first determined that PAE did not change DAPI stained nuclei concentrations within this region across exposure or sex (*Fig.* 4A). Next, we found that quantification of raw mRNA transcript abundance mirrored the patterns of expression observed in co-localization of mRNA with DAPI-stained nuclei (*Supplementary Fig. 3*); therefore, all data are subsequently presented by cell + expression, rather than raw transcript levels. The following comparisons were selected *a priori* for nested analyses of cellular subtypes, with $p$-values corrected for multiple comparisons: 1) Control Males vs Control Females, 2) Control Males vs PAE Males and 3) Control Females vs PAE females.

In quantification of CRFR1+ cells, control males and females did not differ in their proportional expression in CeM cells [$t$(1,67) = 2.029, $p$ = 0.133] (*Fig. 4B)*. However, exposure produced significant changes in % CRFR1+ cells within this region in both males [$t(1,67) = 3.711$, $p = 0.001$] and females [$t(1,67) = 5.966$, $p < 0.001$]. Importantly, this change was bidirectional between sexes, with PAE producing a significant reduction in CRFR1+ cells in males, and a significant increase in CRFR1+ cells in females.

Quantification of total CB+ cells within this region revealed a significant effect of sex in control animals, with control females exhibiting lower proportions of CB+ cells than males [$t(1,67) = 2.767$, $p = 0.022$] (*Fig. 4C).* PAE did not change the proportion of CB+ cells within this region in females [$t(1,67) = 1.220$, $p = 0.538$], however PAE significantly reduced the percentage of CB+ cells in males [$t(1,67) = 2.863$, $p = 0.017$]. In CRFR1+ cells, approximately 1/3 of cells co-labeled with CB in both male and female controls, with no statistical difference between the two groups [$t(1,67) = 0.639$, $p = 0.893$]. PAE did not significantly change the proportion of CRFR1+ cells co-labeled with CB in females [$t(1,67) = 0.670$, $p = 0.879$] or males [$t(1,67) = 2.265$, $p = 0.079$], although males demonstrated a non-significant trend toward a PAE-induced decrease in CRFR1-CB co-labelled cells (*Fig. 4D).*

Quantification of total SST+ cells within this region revealed no difference in proportion of SST+ cells between control males and females [$t(1,67) = 0.345$, $p = 0.980$] (*Fig. 4E).* PAE did not change the proportion of SST+ cells within this region in females [$t(1,67) = 0.730$, $p = 0.849$], however PAE significantly reduced the percentage of SST+ cells in males [$t(1,67) = 2.634$, $p = 0.031$]. In CRFR1+ cells, approximately 30% of cells co-labeled with SST in both male and female controls, with no statistical difference between the two groups [$t(1,67) = 0.233$, $p = 0.994$]. PAE did not significantly change the proportion of CRFR1+ cells co-labeled with SST in females [$t(1,67) = 0.093$, $p = 0.999$] or males [$t(1,67) = 1.883$, $p = 0.232$] (*Fig. 4F).* There was minimal overlap between SST+ and CB+ cells within this region (<5% across experimental groups), and therefore double-labeled SST/CB cells were not statistically analyzed due to insufficient power.

***PAE produces sex-specific changes in behavioral response to CRFR1 agonist infusion in the CeM.*** In our final series of experiments, we evaluated whether our lab's model of moderate PAE changed behavioral response to the activation of CRFR1 within the CeM via a region-targeted infusion of 100 nM Stressin-1. This concentration was selected because it produced exposure-specific physiological response in males, but not females, in our electrophysiology assessments of mIPSC frequency (*Fig. 2)*. We used the LDB assay to assess multiple measures of generalized anxiety-like behavior.

When assessing time spent in the light chamber of the apparatus, a significant interaction between drug infusion (ACSF, Stressin-1) and exposure was uncovered in males [$F(1,33) = 5.800$, $p = 0.022$] (*Fig. 5A),* an interaction that was not statistically significant in females [$F(1,33) = 3.184$, $p = 0.084$]. Although not statistically significant, females demonstrated a trend toward an effect of exposure [$F(1,33) = 4.055$, $p = 0.052$] that was completely absent in males [$F(1,33) = 0.273$, $p = 0.605$], while males demonstrated a non-significant trend toward an effect of drug [$F(1,33) = 3.646$, $p = 0.065$] that was completely absent in females [$F(1,33) = 0.293$, $p = 0.592$]. To further elaborate on sex-specific effects, males were separated graphically by exposure to depict group-specific drug effects (*Figs. 5B,C,G,H)* and females were separated graphically by drug infusion to depict group-specific exposure effects (*Figs. 5D,E,I,J)*. Follow-up analyses of time spent in the light chamber revealed that in males, Stressin-1 infusion in control animals reduced overall time spent in the light chamber [$t(1,33) = 2.867$, $p = 0.014$] (*Fig. 5A&B)* but did not change time spent in the light chamber in PAE males [$t(1,33) = 0.379$, $p = 0.914$] (*Fig. 5C)*. Control males and PAE males did not statistically differ in their time spent in light chamber when infused with ACSF, although there does appear to be a notable reduction in time spent in the light chamber by PAE males [$t(1,33) = 2.051$, $p = 0.094$]. In females, there was no effect of Stressin-1 infusion in either control [$t(1,33) = 0.918$, $p = 0.597$] or PAE females [$t(1,33) = 1.580$, $p = 0.232$], however, PAE females infused with ACSF demonstrated significantly less time in the light chamber than control females infused with ACSF [$t(1,33) = 2.644$, $p = 0.025$] (*Fig. 5A&D)*. This difference was not present between exposure groups following infusion of Stressin-1 [$t(1,33) = 0.165$, $p = 0.983$] (*Fig. 5E)*.

When assessing the number of transitions between light and dark chambers, males once again demonstrated a significant exposure x drug infusion interaction [$F(1,33) = 4.225$, $p = 0.048$] (*Fig. 5F),* and once again this interaction was not significant in females [$F(1,33) = 0.524$, $p = 0.474$]. There were no main effects of drug infusion in either sex (males: [$F(1,33) = 0.332$, $p = 0.569$]; females: [$F(1,33) = 0.226$, $p = 0.637$]) and no main effect of exposure in males [$F(1,33) = 0.010$, $p = 0.922$]. Although not statistically significant, there was a notable trend toward a main effect of exposure in females [$F(1,33) = 3.164$, $p = 0.084$] that appears predominantly driven by PAE, with PAE reducing the number of transitions throughout the test (*Fig. 5I)*. There was no significant effect of exposure in females infused with Stressin-1 [$t(1,17) = 0.791$, $p = 0.386$] (*Fig. 5J*). Post-hoc analyses in males revealed a non-significant pattern a reduction in transitions following infusion of Stressin-1 in control males [$t(1,14) = 3.695$, $p = 0.075$] (*Fig. 5F&G)* with no effect in PAE males [$t(1,13) = 0.933$, $p = 0.346$] (*Fig. F&5H)*.

When examining transitions across time bins, we observed that control males infused with ACSF demonstrated consistent rates of crossing between chambers throughout the testing period. In contrast, control males infused with Stressin-1 did not transition for at least 9 min after initially entering the dark chamber. To determine if transitioning behavior was time-specific, we ran additional post-hoc analyses investigating changes in transition rates between time bins. There was no significant effect of time in control males infused with ACSF ($p = 0.224$), but there was a significant effect of time in control males infused with Stressin-1 ($p = 0.037$) (*Fig. 5G)*.

When assessing time to first enter the dark chamber (egress latency), there were no significant effects of drug infusion [$F(1,30) = 0.068$, $p = 0.069$] or exposure [$F(1,30) = 0.191$, $p = 0.665$] in males, nor a significant drug x exposure interaction [$F(1,30) = 1.927$, $p = 0.176$]. In females, there was also no effect of drug infusion [$F(1,33) = 0.068$, $p = 0.240$], nor an interaction of drug x exposure [$F(1,33) = 3.486$, $p = 0.071$]; however, there was a significant effect of exposure [$F(1,33) = 4.515$, $p = 0.041$]. Post-hoc analyses determined this significant effect of exposure was once again specific to ACSF-infused females, with PAE reducing egress latency in this group [$t(1,33) = 2.779$, $p = 0.018$], while exposure did not change response to Stressin-1 infusion in females [$t(1,33) = 0.185$, $p = 0.979$].

Assessments of time to first return to the dark chamber (re-entry latency) revealed no significant effects of drug infusion [$F(1,31) = 1.310$, $p = 0.261$], exposure [$F(1,31) = 0.129$, $p = 0.722$] or an interaction of these variables [$F(1,31) = 2.567$, $p = 0.119$] in males. Similarly, these factors did not influence re-entry latency in females: (drug infusion: [$F(1,33) = 0.455$, $p = 0.504$]; exposure: [$F(1,33) = 2.621$, $p = 0.115$]; drug infusion x exposure interaction: [$F(1,33) = 0.423$, $p = 0.520$]. Assessment of head poking from the dark chamber into the light chamber produced similar null results: no significant effects of drug infusion [$F(1,32) = 1.844$, $p = 0.184$]. exposure [$F(1,32) = 0.717$, $p = 0.407$] or an interaction of these variables [$F(1,32) = 2.354$, $p = 0.135$] in males, and no significant effects of drug infusion [$F(1,33) = 0.001$, $p = 0.973$], exposure [$F(1,33) = 0.022$, $p = 0.883$] or an interaction of these variables [$F(1,33) = 3.268$, $p = 0.077$] in females.

*Supplementary Material References*
